# Meiotic cohesin paralogs govern cell survival by exhibiting flexibility in partner choice

**DOI:** 10.64898/2026.03.27.714714

**Authors:** Sayantan Banerjee, Purva, Ananya Dodamani, Akriti Kumari, Sudipto Singha, Oankar Varde, Gunjan Mehta, Mridula Nambiar

## Abstract

Erroneous loading of the ring-shaped cohesin complex, especially at centromeres, cause chromosomal segregation defects in both mitosis and meiosis. Mitotic cohesin subunits of this complex, either get replaced or co-exist with their meiotic paralogs during meiosis and also in certain cancers. However, it is unclear whether meiotic paralogs can partner mitotic subunits to form hybrid complexes in somatic cells and if there are any functional consequences on cancer progression. Here, we provide a conceptual framework for the principles of cohesin complex assembly involving non-canonical subunits in proliferating *Schizosaccharomyces pombe*. We show that chromosome loading, segregation fidelity and cellular proliferation are critically affected by the composition of the available cohesin complexes. We find stark differences in the ability of the meiotic kleisin subunit Rec8 to support robust centromere loading, irrespective of its partner, when compared to the canonical mitotic paralog Rad21. Such variations in cellular growth can be explained by different dwell times of these cohesin complexes on the chromosomes as determined by single-molecule tracking and altered chromatin enrichment. We also discover a unique feature of Rec8, in stabilizing chromatin-bound hypomorphic cohesin mutants that aid in cell survival under restrictive conditions. Overall, we highlight the flexibility of meiotic cohesins in restoring function, albeit at a fitness cost, in the presence of inactivating cohesin mutations. Such imbalances could be exploited by cancers to aid cell survival, but at the expense of increased aneuploidy and genomic instability.

## Introduction

Cohesin complexes are canonically made up of four core subunits that include two structural maintenance of chromosomes (SMC) subunits, SMC1 and SMC3, the kleisin RAD21 and the HEAT repeat protein associated with Kleisin (HAWK), STAG1 and STAG2. In *Schizosaccharomyces pombe,* the orthologs of these proteins are Psm1, Psm3, Rad21 and Psc3, respectively. Deletion of any one of the subunits results in lethality, making all of them essential for cell survival. Interestingly, the kleisin Rad21 and HAWK subunit Psc3 have meiosis-specific paralogs, Rec8 and Rec11, whose ectopic expression can rescue cell inviability in their absence in proliferating cells^1,2^. During meiosis, Rec8-Rec11 complex is deposited at the chromosomal arms whereas Rec8-Psc3 is enriched at the centromeres^1^. These spatially separated cohesin populations are necessary to prevent activation of deleterious meiotic DSBs and recombination at the centromeres that can cause aneuploidy in gametes^2,3^. In human and mouse somatic cells, there are two STAG proteins that perform locus-specific functions on chromosomes – STAG1 maintains telomere cohesion, while STAG2 is required for centromere cohesion^4,5^. Germline cells have additional meiosis-specific kleisins (REC8 and RAD21L1) and HAWK protein STAG3, that further add to the pool of cohesin complexes available for chromosomal loading^6–8^. Interestingly, aberrant expression of meiotic cohesins such as REC8, STAG3 and SMC1β has been reported in various types of cancers, which can lead to simultaneous existence of multiple potential cohesin complexes that may or may not have a functional consequence, unless loaded on the chromosomes^9–11^. It is unclear whether these non-canonical hybrid complexes between mitotic and meiotic paralogs are formed at all and if they have any effect on cellular growth. Hence there is a need to systematically address the potential functional roles and effects of having these different subtypes of cohesins in cells under different conditions.

Targeting and loading of cohesins at the heterochromatin flanking the centromeres (pericentromere) is thought to be largely facilitated via the abundant heterochromatin protein 1 (HP1) ortholog Swi6 that binds to the H3K9 methylated histones via its chromodomain (CD) and can additionally interact with the cohesin loader complex Mis4-Ssl3 in *S. pombe*^12–14^. Additionally, Swi6 can directly interact with the HAWK cohesin subunit Psc3 and along with Hsk1-Dfp1 DDK kinase activity further facilitates cohesin loading^12,15^. In absence of Swi6, cells exhibit increased chromosomal segregation errors during both mitosis and meiosis leading to fitness loss and reduced meiotic spore viability, due to substantial loss of pericentromeric cohesion^12,14^. These observations argue majorly for a Psc3 and Swi6 driven cohesin loading at the heterochromatin that may have additionally evolved to actively exclude the recombination-activating meiotic Rec11 cohesin complex from the pericentromeric heterochromatin during meiosis^2,3,16^. Furthermore, Rec8-Psc3 cohesin complex gets loaded at the central core of the centromere to aid in reductional division of meiotic chromosomes, although the factors involved are unknown^1^. In *S. cerevisiae,* cohesin loading at the centromere core is mediated by the inner kinetochore protein Ctf19 with the help of DDK kinase activity^17^. In the chromosomal arms, chromatin remodelers such as the Rsc complex have been implicated in facilitating cohesin loading via interaction with the loader subunit Ssl3^18–20^. Although we largely know the molecular players involved in cohesin loading, there is very limited understanding of how these factors come together in dictating the specificities of which population to deposit across genomic loci, when there are multiple cohesin complexes to choose from. Mechanistically, this should primarily be driven by principles of cohesin complex formation, based on the availability of the type of cohesin subunits in the cell. It is unknown if there are any inherent rules for formation and functional loading of cohesin complexes on the chromosomes in the presence of multiple kleisin and HAWK paralogs that may affect chromosomal segregation and viability. Deciphering the consequences of disrupting any such preferences among the cohesin paralogs may help understand the origin of both mitotic and meiotic aneuploidies in cancers and developmental disorders, respectively.

We started by asking whether meiotic cohesin subunits can form non-canonical complexes with their mitotic partners and are these cohesins functional in proliferating *S. pombe* cells. We systematically generated a pool of cohesin complexes containing different combinations of kleisin and HAWK paralogs, mimicking meiotic cells or cancer cell-like states that aberrantly express meiosis-specific cohesins. We find that similar to meiosis, at the mitotic centromeres, the meiotic HAWK Rec11 cohesins are excluded, but can be overcome by direct targeting of the complex to the pericentromeric heterochromatin. We provide evidence that the composition of the cohesin complex at the centromeres dictates their chromatin dwell times, which can be correlated to the chromosomal segregation fidelity in these cells. We also discovered a distinctive property of Rec8 in stabilizing chromatin-bound Psc3 hypomorphic mutants, which could restore cell growth even at non-permissive conditions, although at a fitness cost. Overall, our study demonstrates that presence of meiotic or hybrid cohesin complexes can aid cellular proliferation, when canonical cohesin loading mechanisms are impaired. This flexibility can be exploited as a valuable strategy by cancer cells that frequently harbor inactivating cohesin mutations and aberrantly express meiotic paralogs, to escape complete loss of cohesin function.

## Results

### Meiotic cohesin Rec8 supports cellular growth irrespective of HAWK partner unlike mitotic paralog Rad21

In order to determine the extent to which the meiotic cohesins can complement the loss of canonical Psc3-Swi6 driven cohesin loading, we utilized three different temperature-sensitive alleles of Psc3 (referred to as -*2ts, -3ts, -4ts* alleles) that have been previously reported^12^. We sequenced these three alleles and found that each possesses a different set of unique point mutations (Suppl. Fig. 1A). Each *psc3-ts* allele was completely sensitive at the non-permissive 34°C with *psc3-4ts* being the most unstable (Suppl. Fig. 1B). In the absence of Psc3, the assumption is that cells undergo massive chromosomal segregation errors due to absence of cohesion, especially at the pericentromeric heterochromatin, thereby leading to cell death^12,14,15^. Since the meiotic paralog Rec11 forms functional cohesin complexes at chromosomal arms in meiosis, we tested whether Rec11 could functionally complement Psc3 in proliferating cells. However, when *rec11* was ectopically expressed under a constitutive promoter (*rec11ee)* in *psc3-4ts* cells, we observed no rescue in growth at the non-permissive temperature, unless the meiotic kleisin Rec8 was also constitutively co-expressed (*rec8ee)* (Fig. 1A). This was true for the other two mutant alleles *psc3-2ts* and *psc3-3ts* as well (Suppl. Fig. 1C). This is also supported by inability to obtain a *psc3Δ rec11ee* genotype during genetic crosses or direct *psc3* deletions via transformations, unless *rec8ee* is concomitantly present (Suppl. Fig. 1D). Additionally, we also observed that all three *psc3-ts* mutants were hypomorphs, since they showed varying growth rates even at the permissive temperature of 25 °C, which stayed unaffected in presence of *rec11ee* (Fig. 1B and Suppl. Fig. 1E). Interestingly, additional expression of *rec8,* restored the growth rates to that seen for *rec11ee* alone (Fig. 1B and Suppl. Fig. 1E). This result highlights that growth defects in presence of Psc3 hypomorphic mutants can be restored by concomitant expression of meiotic cohesins Rec8 and Rec11.

**Figure 1.**
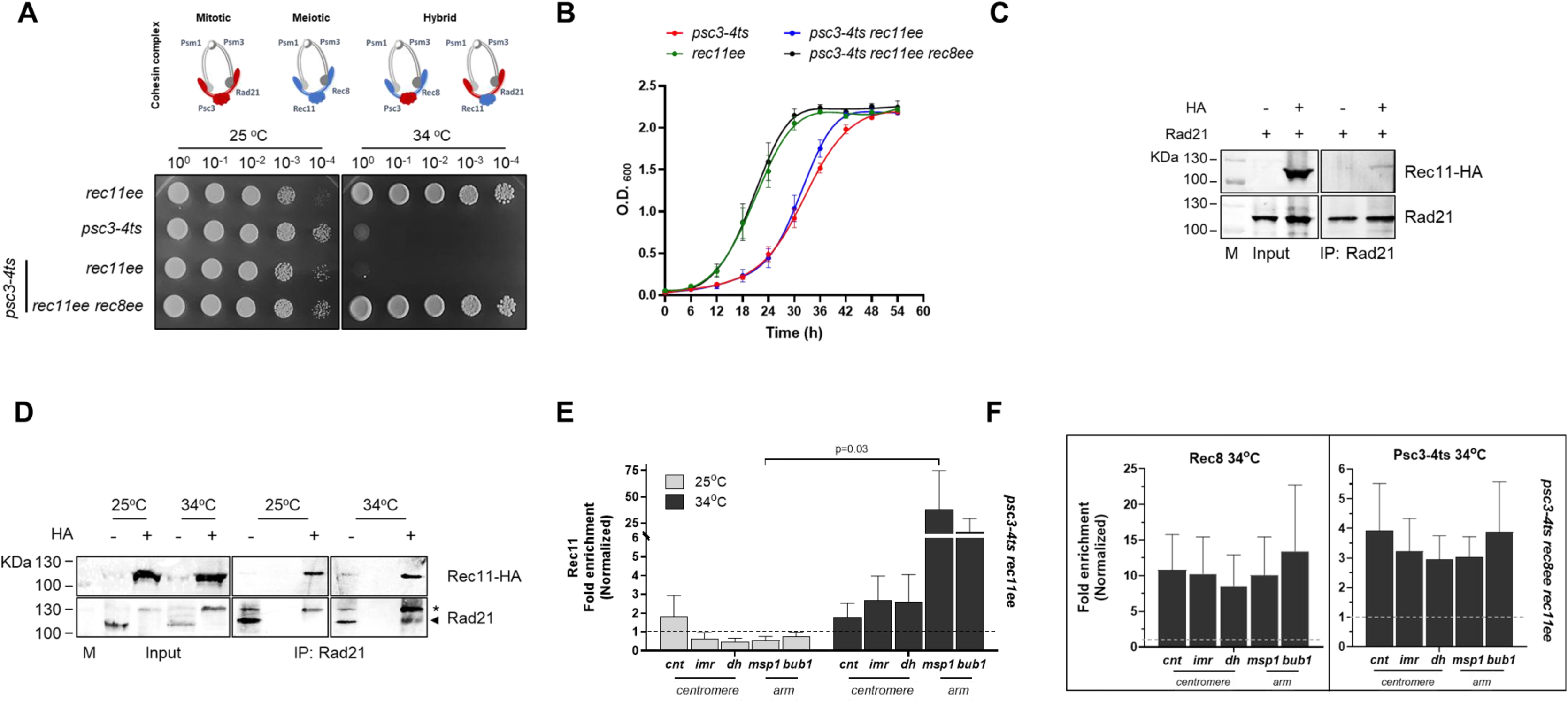
The canonical meiotic cohesin complex Rec8-Rec11 rescues viability in the absence of mitotic HAWK subunit Psc3. **A.** Spot assay for *psc3-4ts* strains in the presence of ectopic expression (ee) of its paralog Rec11 (*rec11ee)* alone or with its meiotic partner Rec8 (*rec8ee)* at 25 °C and 34 °C. **B.** Growth curves for the strains in panel A at 25 °C. **C-D.** Interaction between Rad21 and Rec11 in wild type (C) and *psc3-4ts* (D) cells with and without overexpression of *rec11-HA* via co-immunoprecipitation. Rad21 was immunoprecipitated and probed with anti-HA and anti-Rad21 antibodies. M is ladder. Arrowhead shows the Rad21 band seen in wild type cells, whereas the asterisk marks Rad21 seen in the presence of Rec11 expression, as seen in the respective inputs. **E-F.** ChIP qPCR for Rec11 (E) and Rec8 and Psc3-4ts (F) in *psc3-4ts rec11-HAee* and *psc3-4ts rec8ee rec11-HAee*, respectively, at the indicated temperatures. Both centromeric (*cnt, imr, dh*) and arm (*msp1, bub1)* loci were tested, wherein *cnt* is from the centromere core region, *imr* from the innermost repeat, *dh* is at one of the outer repeat regions, *msp1* is a cohesin-rich arm region and *bub1* is a control arm region. Fold enrichment was calculated after normalizing with mitochondrial DNA locus as negative control. For each experiment, atleast 3 biological repeats were performed. For panels B, E and F, bars represent the mean of at least three independent experiments performed in triplicates; error bars=SEM.

Since the Psc3 hypomorphic mutants also natively express Rad21, and addition of Rec11 alone was unable to rescue viability, it indicates two possibilities. Either there is incompatibility between Rec11 and Rad21 to form a physical complex or if formed, the complex may not sufficiently get loaded on the chromosomes, thereby affecting vital processes including chromosomal segregation. We tested the formation of protein complex between Rad21 and ectopically expressed Rec11-HA in wild type cells via co-immunoprecipitation (co-IP) using chromatin fraction and observed a weak but reproducible interaction (Fig. 1C and Suppl. Fig. 1F). We also confirmed the interaction between Rad21 and Rec11 in a *psc3-ts* background at both the permissive and non-permissive temperatures (Fig. 1D). In these cells, we observed migration of Rad21 at a higher MW when Rec11 was expressed in the cells, possibly suggesting presence of a differentially modified form of Rad21. Additionally, we also tried to predict the interaction between the kleisin subunits and Rec11 using Alphafold 3 (Suppl. Fig. 2A, 2B). Rad21 was found to have a largely disordered structure with three main domains, viz., N-terminal, C-terminal domains which interacts with the Psm3 and Psm1 subunits, respectively and a middle domain, whose function remains yet to be attributed (Suppl. Fig. 2A). The middle domain was separately tested and found to interact with Rec11/Psc3. We have named this as the domain for HAWK interaction (DHI). AlphaFold predicted a high confidence interaction between Rad21^DHI^ and Psc3 (iPTM > 0.7), while it failed to confidently predict any Rec11-Rad21^DHI^ interaction (iPTM ∼ 0.54) (Suppl. Fig. 2B). This low confidence interaction along with our co-IP data point towards weak interaction between Rad21 and Rec11, which may affect the stability and the fraction of functional cohesins available for chromatin loading in cells.

Since, the Rad21-Rec11 complex could form, although not robustly, we wondered if the cells were inviable due to insufficient chromatin-bound cohesin levels. We tested for the enrichment levels of Rec11 across multiple loci within the centromeres and the chromosomal arms via ChIP quantitative PCR. There was no significant Rec11 enrichment at any of the centromeric loci at either the permissive (25 °C) or non-permissive (34 °C) conditions for Psc3 depletion (Fig. 1E). One of the arm loci showed significant enrichment of Rec11, indicating minimal cohesin loading on the euchromatin. Hence, cell inviability could be attributed to the overall lack of cohesion leading to segregation defects.

We further confirmed that the rescue of growth defects at 34 °C in the presence of both Rec8 and Rec11 as seen above, could be explained by restoration of cohesin loading at different genomic loci, especially the centromeres (Fig. 1F, 1A). Surprisingly, enrichment of Psc3-4ts mutant protein was also observed at 34 °C, when Rec8 was present, which could contribute towards the rescue (Fig. 1F, 1A). This suggests possible persistence of the mutant Psc3 protein even at the restrictive 34 °C, which we will further discuss separately below. Overall, these data strongly suggest that despite availability of cohesin complexes they may not always be functional and show robust chromatin-binding. The hybrid Rad21-Rec11 cohesin fails to restore centromeric cohesion, unlike the meiotic Rec8-Rec11, showing partner choice in cohesin complexes playing a key role in cellular proliferation.

### Targeting the non-canonical Rad21-Rec11 cohesin complex to the pericentromeric heterochromatin is sufficient to restore cellular growth

The inability to rescue inviability upon depletion of Psc3 by Rad21-Rec11 complex, could plausibly be due to insufficient centromeric cohesion, leading to chromosomal missegregation (Fig. 1E). Hence, we tested whether restoration of cohesins at the pericentromeric heterochromatin by artificially tethering Rad21-Rec11 complex via Rec11-Chromodomain (CD) fusion (*rec11-CD*) could rescue viability by binding to the H3K9me histones at the heterochromatin. Indeed, we were able to now observe a complete rescue in cell viability, upon inactivation of Psc3 (*psc3-4ts* and *psc3-2ts* alleles) at the non-permissive 34 °C, but only when Swi6 was additionally removed (Fig. 2A and Suppl. Fig. 2C). This suggested that upon depletion of Psc3, the alternative Rad21-Rec11 complex could not get deposited at the heterochromatin (in the presence of Swi6) on its own. We further confirmed significant Rad21 cohesin enrichment at the centromeres at both 25 °C and 34 °C, but only when Swi6 was removed, supporting the cell survival seen above (Figs. 2A, 2B). Interestingly, we were also able to capture weak interaction between Rec11 and Swi6, which suggests the possibility of basal, albeit insufficient, chromatin loading of Rec11 cohesin complexes at the heterochromatin (Suppl. Fig. 2D). These data provide evidence that restoration of adequate centromeric cohesion, irrespective of the type of cohesin complex loaded, is sufficient to atleast allow cell proliferation and survival in the absence of the canonical cohesin loading mechanisms.

**Figure 2.**
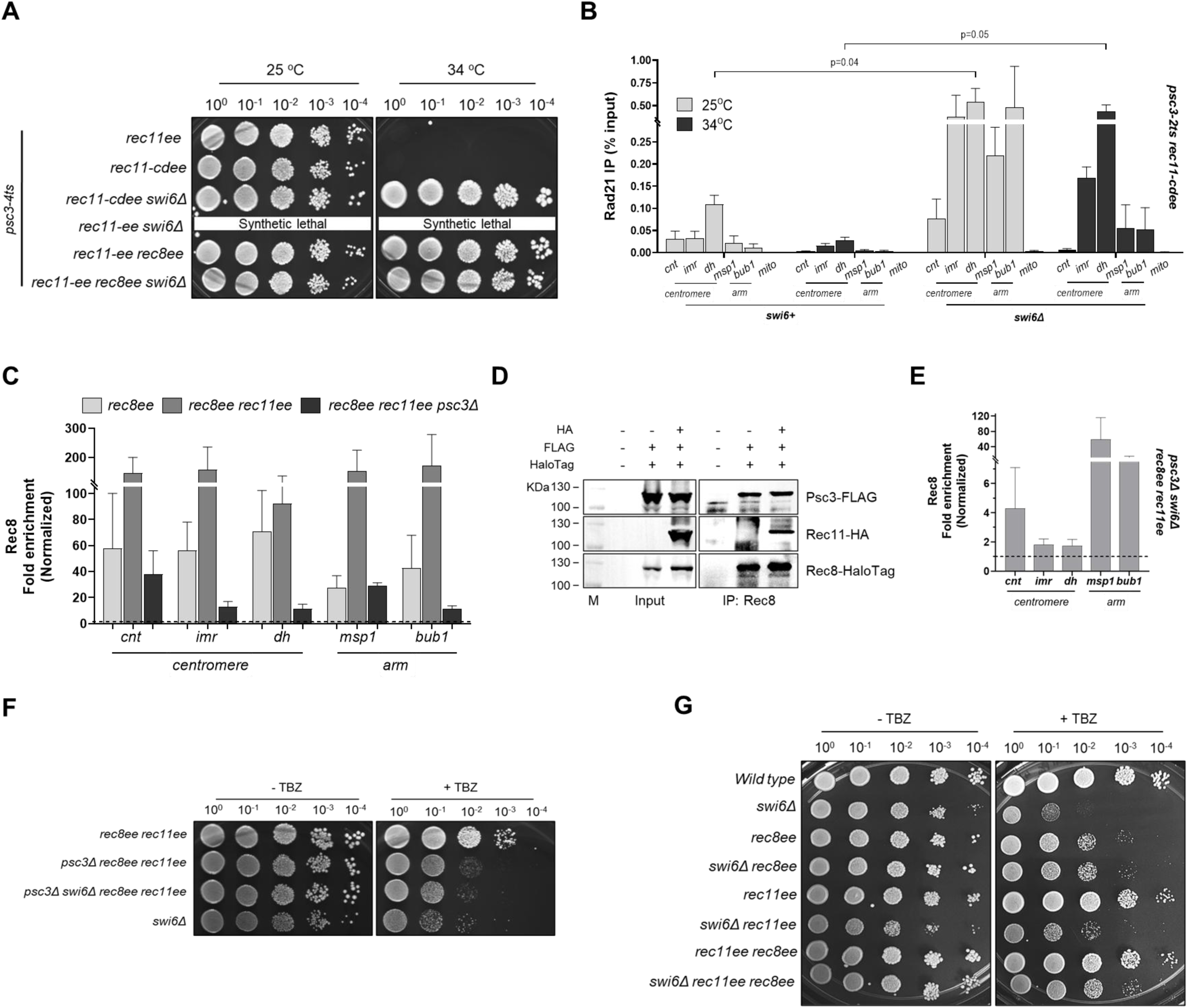
The meiotic kleisin Rec8 facilitates cohesin loading with Rec11 at centromeres unlike mitotic paralog Rad21. **A.** Spot assay for *psc3-4ts* strains in the presence of *rec11ee, rec11-cdee* (Rec11 fused with Swi6 chromodomain (CD))*, rec8ee* with and without *swi6Δ* at 25 °C and 34 °C. **B.** ChIP qPCR for Rad21 in *psc3-2ts rec11-cdee swi6+* and *swi6Δ,* at the indicated temperatures. Percent input was calculated for the Rad21 enrichment. **C.** ChIP qPCR for Rec8-HA in *rec8ee, rec8ee rec11ee* and *rec8ee rec11ee psc3Δ*. Fold enrichment was calculated after normalizing with mitochondrial DNA locus as negative control. **D.** Interaction between Rec8, Psc3 and Rec11 in wild type cells overexpressing both Rec8 and Rec11 via co-immunoprecipitation. Rec8-HaloTag was immunoprecipitated and probed with anti-HA (for Rec11) and anti-FLAG (for Psc3) antibodies. M is ladder. **E.** ChIP qPCR for Rec8-HA in *psc3Δ swi6Δ rec8ee rec11ee*. Fold enrichment was calculated after normalizing with mitochondrial DNA locus as negative control. **F-G.** Spot assay to assess hypersensitivity to Thiabendazole (TBZ) (0 and 7.5 μg/ml) in *psc3Δ* and *psc3Δ swi6Δ* (F) and *swi6Δ* (G) backgrounds upon expression of Rec8 and/or Rec11. For each experiment, atleast 3 biological repeats were performed. For panels B, C and E, bars represent the mean of at least three independent experiments performed in triplicates; error bars=SEM.

### Rec8 bypasses canonical preferences of centromeric cohesin loading and allows flexibility in cell survival

We observe that between the two kleisin subunits Rad21 and Rec8, the meiotic paralog provided more flexibility in terms of ability to be chromatin-bound upon depletion of Psc3 (Figs. 1A, 1F). In order to test, if the Rec8 subunit was utilized for complex formation and functional complementation, only when the canonical Rad21-Psc3 complex was limiting, we determined the chromatin-bound levels of Rec8 in various contexts, in the presence of Rad21. We tested for chromatin-bound Rec8 levels in *rec8ee* (enriching for Rec8-Psc3), *rec8ee rec11ee* (enriching for both Rec8-Psc3 and Rec8-Rec11) and *rec8ee rec11ee psc3Δ* (enriching for Rec8-Rec11) across multiple chromosomal loci and found it to be enriched at all tested loci, including the centromere core, as has been reported previously^21,22^ (Fig. 2C). When Rec11 was co-expressed, Rec8 occupancy appeared to further increase across all loci, especially in the chromosomal arms, mimicking the scenario in meiosis, where also two types of cohesin complexes are available (Fig. 2C). We confirmed the interaction of Rec8 with both Psc3 and Rec11, when expressed together, demonstrating the formation of both the complexes (Fig. 2D). When *psc3Δ* was introduced in the *rec8ee rec11ee* background, although there was a drop in the fold enrichment, the Rec8 occupancy continued to be high both at the centromeres and the arms, clearly pointing towards loading of the Rec8-Rec11 complex (Fig. 2C). These results highlight the co-existence of multiple type of cohesin complexes on the chromatin and the innate flexible property of the meiotic kleisin Rec8 to provide robust alternative options that its mitotic counterpart Rad21 fails to offer.

We further went on to test whether the Rec8-Rec11 complex could be functionally utilized to bypass the synthetic lethality of Psc3 and Swi6, as has been reported earlier, probably due to failure of cohesin loading^12^. Indeed, we found enrichment of Rec8 in *psc3Δ swi6Δ* background, although the occupancy at the pericentric loci (*imr* and *dh*) was low compared to that at the centromere core (*cnt)* and chromosomal arms (Fig. 2E). Cohesin loading at the centromere core regions is a hallmark of meiotic chromosomes to facilitate mono-orientation and is driven by Rec8^23,24^. Hence, our data clearly show that the enrichment of cohesins at the centromere core, along with that in the chromosomal arms, is sufficient to support growth and can be utilized for cell survival as a backup mechanism.

### Composition of the centromeric cohesins dictates fidelity of chromosomal segregation

Usage of alternative meiotic cohesin complexes was sufficient to restore cell viability as seen above, but are there associated fitness costs? We measured hypersensitivity to the spindle poison Thiabendazole (TBZ), in *psc3Δ swi6Δ rec8ee rec11ee* cells, with disrupted canonical cohesin loading at the pericentromeric heterochromatin, in order to assess the fidelity of chromosomal segregation. Although ectopic overexpression of both Rec8 and Rec11 does not induce much errors on its own in a wild type background, absence of Psc3 and/or Swi6 makes the cells more prone to mis-segregation (Fig. 2F).

Swi6 is a key regulator of cohesin loading at the pericentromeric heterochromatin and its absence makes the cells highly sensitive to TBZ. Interestingly, these defects were largely rescued by ectopic expression of *rec8* independent of *rec11,* despite presence of Rad21 (Fig. 2G). Since *rec8ee* alone showed some sensitivity to TBZ on its own, the extent of rescue in *swi6Δ* upon expression of Rec8 and/or Rec11 was similar to that in *rec8ee* alone and did not reach wild type levels (Fig. 2G). Overall, it appears that the canonical Rad21-Psc3 complex assists in the most precise chromosomal segregation in proliferating cells, whereas the meiotic Rec8-Rec11 is error-prone when Psc3 is absent, due to low cohesion at the pericentromeres (Figs. 2E, 2F). However, in a wild type background (*rec8ee*), presence of Rec8-Psc3 induces mis-segregation, probably due to the presence of cohesins at the centromeric core regions, as has been implicated before^21,25^ (Fig. 2C). Interestingly, in this context, co-expression of Rec11 (*rec8ee rec11ee*) can actually rescue some of these defects and restore segregation fidelity (Fig. 2G). These observations clearly show that although utilizing alternate available meiotic cohesin complexes can be advantageous to the cells in terms of survival, it definitely comes at a cost.

### Rec8-Psc3 and Rec8-Rec11 cohesins have different dwell times on chromatin

To understand why meiotic cohesins are insufficient in supporting faithful chromosomal segregation during mitosis, we wondered if complexes with varying compositions have differences in their binding and stability on chromatin. For this, we performed single-molecule tracking of Rec8-HaloTag in *rec8ee* (having only Rec8-Psc3 complex), *rec8ee rec11ee* (having both Rec8-Psc3 and Rec8-Rec11 complexes) and *rec8ee rec11ee psc3Δ* (having only Rec8-Rec11 complex) to quantify their chromatin dwell times and bound fractions. Rad21 was present in all the genotypes, hence we assume its competitive effect should be similar across all the genetic backgrounds. We tracked Rec8-HaloTag on heterochromatin (marked by Swi6-GFP) and euchromatin separately (Fig. 3A). The dwell time distribution fit double exponential decay curve, suggesting two types of bound molecules of Rec8-HaloTag - 1) bound with long dwell time (pink fraction), and 2) bound with short dwell time (blue fraction) (Fig. 3B). Previous work suggests that the pink fraction represents specific, bound molecules that would have a biological function, whereas the blue fraction represents non-specifically bound molecules that make non-specific, transient interactions with the chromatin without any function^26–29^. Interestingly, we observed differences in the dwell times and specific bound fractions (pink fractions) of all these complexes between heterochromatin and euchromatin (Fig. 3B, 3C). The Rec8-Psc3 complex showed more stable interaction with the heterochromatin, when compared to euchromatin, as its dwell time increased from 4.97±0.2 s at euchromatin to 19.1±2.4 s at heterochromatin, with a modest reduction in the specific bound fraction at heterochromatin (from 33.3±1.6% at euchromatin to 27.9±2.4% at heterochromatin) (Fig. 3B, 3C). This result suggests that the stability of Rec8-Psc3 complexes on chromatin may depend on the open or closed architecture of the chromatin.

**Figure 3.**
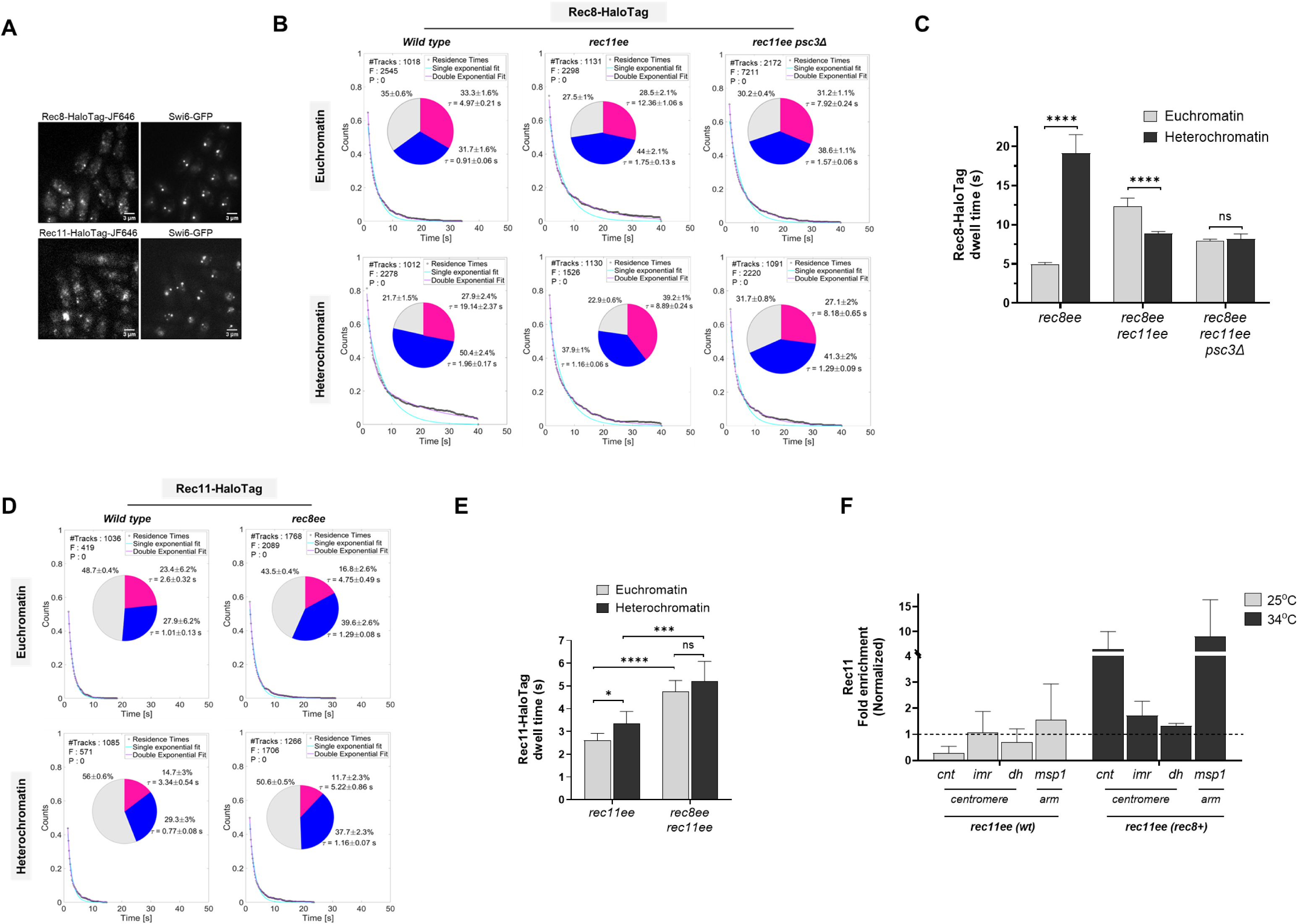
Single molecule tracking of Rec8 and Rec11 to quantify their chromatin binding dwell time and fraction show differences based on the composition of the complex. **A.** Representative images show single molecules of Rec8-HaloTag-JF646 and Rec11-HaloTag-JF646 in the nuclei of cells expressing Swi6-GFP to mark heterochromatin. Scale bar: 3 µm. **B.** Dwell time analysis of Rec8 at euchromatin and heterochromatin using a slow-imaging regime. Survival probability distributions were fitted with single (cyan) or double (magenta) exponential decay curves. For all conditions, the survival distribution fitted best to the double-exponential decay, suggesting two populations of dwell times, represented by the pie charts. The pie charts represent the fraction of molecules bound with long dwell time (pink fraction) and the fraction of molecules bound with short dwell time (blue fraction), along with their mean dwell time (*τ*). The gray fraction represents diffusing molecules. ‘#Tracks’ represents the total number of tracks analyzed, ‘F’ represents the value of the ‘*F* test’ to distinguish between single-versus double-exponential fitting, and ‘P’ represents the p-value of the *F* test. **C.** Bar graph comparing the residence times of Rec8 for *rec8ee, rec8ee rec11ee* and *rec8ee rec11ee psc3Δ* cells, taken from panel B. **D.** Dwell time analysis of Rec11 at euchromatin and heterochromatin using a slow-imaging regime. Survival probability distributions, pie charts and other annotations as described in panel B. **E.** Bar graph comparing the residence times of Rec11 for *rec11ee* and *rec11ee rec8ee* cells, taken from panel D. **F.** ChIP qPCR for Rec11-HA in *rec11ee* alone *(wt)* and *rec11ee rec8ee (rec8+)*. Fold enrichment was calculated after normalizing with mitochondrial DNA locus as negative control. Data in panels B and D were analysed by F test. ****p <0.0001; ***p <0.001; *p <0.05; ns p >0.05. The error bars in panels C, E and F represent ±SEM.

On the other hand, the Rec8-Rec11 complex when present alone (in *psc3Δ* background) didn’t show significant differences in their dwell times and specific bound fractions between euchromatin and heterochromatin (dwell time of 7.9±0.2 s at euchromatin and 8.2±0.7 s at heterochromatin; specific bound fraction of 31.2±1.1 % at euchromatin and 27.1±2%, p<0.05), suggesting its stability on chromatin is unaffected by the open or closed architecture of the chromatin (Fig. 3B, 3C). Therefore, we can infer that Rec8-Psc3 with higher residence times and hence more stable binding on the heterochromatin might result in better pericentromeric cohesion than Rec8-Rec11 alone and thereby result in better segregation fidelity during cell division (Fig. 2F, 2G). The stability could be imparted by the direct Swi6-Psc3 interaction on the heterochromatin^12^. Interestingly, in the presence of both the complexes, Rec8-Psc3 and Rec8-Rec11 (in *rec8ee rec11ee* background), the dwell times of Rec8-HaloTag increased significantly at the euchromatin (from 4.97±0.2 s to 12.4±1.1 s, p<0.001) and reduced significantly at the heterochromatin (from 19.1±2.4 s to 8.9±0.2 s, p <0.001), suggesting that the stoichiometric alterations among the binding partners of Rec8 alters the overall stability of the cohesin complexes on chromatin (Fig. 3B, 3C).

We similarly tracked single molecules of Rec11-HaloTag over the heterochromatin and euchromatin both in the absence and presence of ectopic expression of Rec8 in proliferating cells (Fig. 3A, 3D). We observed a small but significant increase in the dwell time of Rec11-HaloTag on the heterochromatin compared to euchromatin (3.34±0.5 s vs 2.6±0.3 s) in *rec11ee* genotype (Figs. 3D, 3E). The percent bound fraction on the euchromatin was higher than the heterochromatin (23.4±0.6 % vs 14.7±3 %; p<0.05), suggesting that open chromatin facilitates binding of Rec11-Rad21 complex (Fig. 3D). Interestingly, upon Rec8 ectopic expression, although the specific bound fraction Rec11-cohesin complexes remained the same (16.8±2.6% at euchromatin and 11.7±2.3% at heterochromatin, p>0.05; ns compared to wild-type), there was a little but statistically significant increase in the residence times of the Rec11-Rec8 complex both at the heterochromatin and euchromatin (Fig. 3D, 3E). We also confirmed increased occupancy of Rec11 in the presence of Rec8 via quantitative ChIP by PCR (Fig. 3F). Interestingly, increased occupancy was observed mostly at the centromere core and the arm locus, when Rec8 was present (Fig. 3F). Overall, these data suggest that Rec8 complexes are more readily bound on the chromatin, irrespective of the partner HAWK protein, whereas the meiotic Rec11, binds stably only when its canonical partner Rec8 is present.

### Kleisin paralogs Rec8 and Rad21 impart differential response to Psc3 loss of function

As observed above, the presence of Rec8 and Rec11, improved the overall survival and growth of Psc3 hypomorphic mutants (Fig. 1A, 1B and Suppl. Fig. 1C, 1E). Since, it is established that Rec8-Rec11 together are essential to rescue *psc3Δ,* we wondered whether a similar requirement exists to support the growth of Psc3 hypomorphic mutants. We first tested growth at the restrictive 34 °C of *psc3-ts rec8ee* cells without additional expression of the meiotic Rec11 and surprisingly, observed a near complete rescue of growth in all three *psc3-ts* mutants when only Rec8 was expressed (Fig. 4A and Suppl. Fig. 3B). This was unexpected as this shows that Rec8 can somehow negate the loss of functionality of the Psc3 mutants at the restrictive temperatures. Despite complete survivability, these cells grew only slightly better than the Psc3 hypomorphic mutant itself at 25 °C, but continued to show improved doubling time, when Rec11 was co-expressed (Figs. 4B and 1B). Upon further *swi6Δ*, the cell viability was visibly reduced in all three *psc3-ts rec8ee* strains but still the rescue was quite significant (Fig. 4A and Suppl. Fig. 3B). Cells expressing Rec11 additionally appeared to grow robustly with no effect of removal of Swi6, as has been observed above (Fig. 4A and Suppl. Fig. 3B).

**Figure 4.**
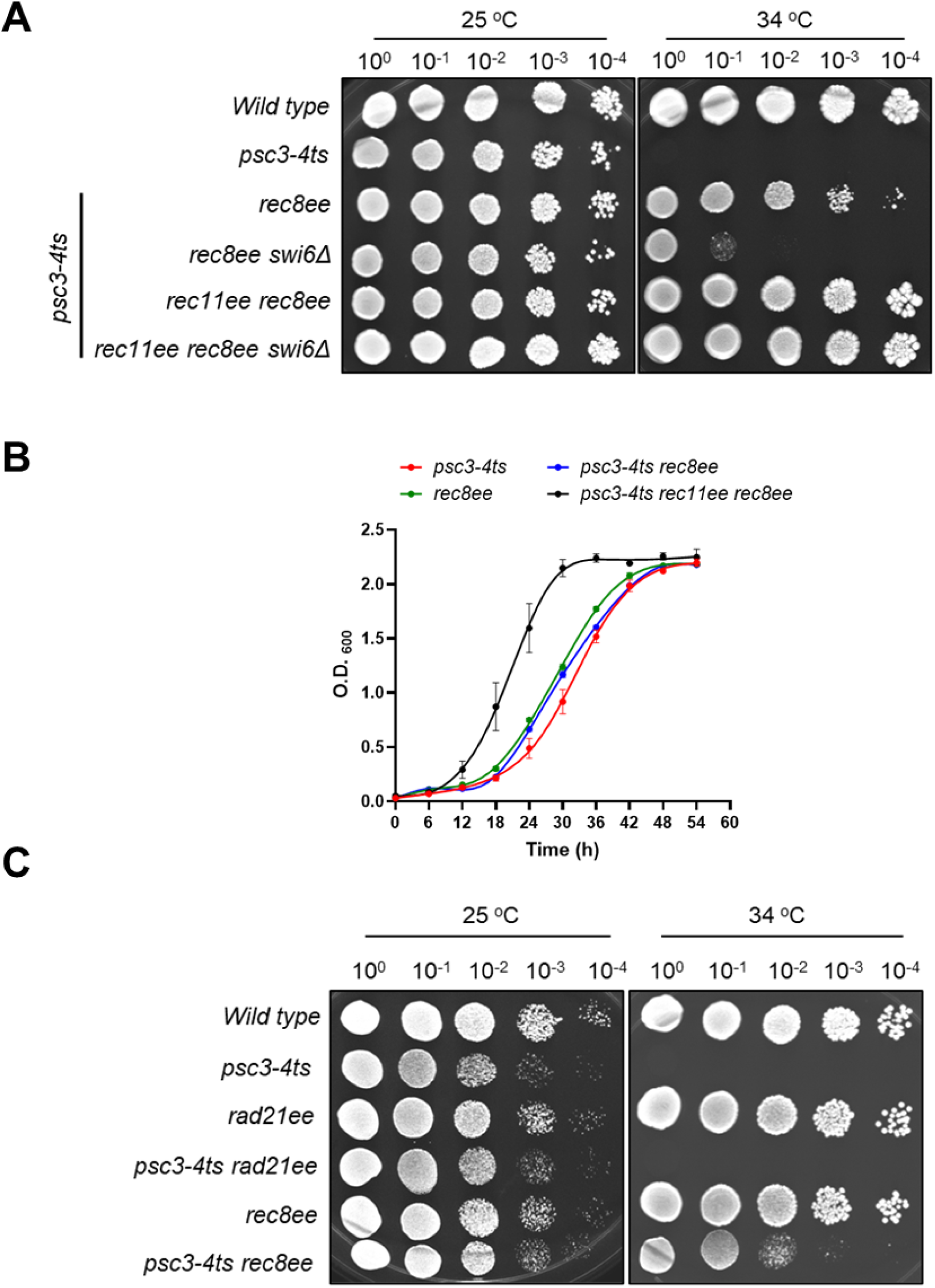
Mitotic and meiotic kleisins show different rescue phenotypes upon interaction with Psc3 hypomorphic mutants under restrictive conditions. **A.** Spot assay for *psc3-4ts* strains in the presence of *rec8ee* with and without *rec11ee* and/or *swi6Δ* at 25 °C and 34 °C. B. Growth curves for *psc3-4ts, rec8ee, psc3-4ts rec8ee* and *psc3-4ts rec8ee rec11ee* at 25 °C. Error bars represent ±SEM. **C.** Spot assay for *psc3-4ts* strains in the presence of *rad21ee* or *rec8ee* at 25 °C and 34 °C. For each experiment, atleast 3 biological repeats were performed.

Since, the natively expressed Rad21 present in these cells is unable to show the same rescue as Rec8, we wondered whether the overexpression of Rec8 was responsible for the rescue, simply due to more complex formation. To test this, we overexpressed *rad21* from the constitutive *padh1* promoter, similar to *rec8.* However, rescue of *psc3-ts* mutants was not observed in the presence of *rad21*ee, confirming it to be a unique and novel property of Rec8 (Fig. 4C and Suppl. Fig. 3C). We additionally looked at the binding between Rec8 and Psc3 using Alphafold 3. Similar to Rad21, Alphafold predicted a similar disordered structure of Rec8 with an unattributed middle DHI domain that was found to interact with Psc3 at a higher confidence score (iPTM >0.7) (Suppl. Fig. 3A). According to the predicted complex, Rad21-DHI has a more disordered structure and interacts with residues of alpha helices on the concave side of Psc3, however, Rec8-DHI has a more helical structure and binds to Psc3 helices more loosely (Suppl. Fig. 3A). We also did not find any of the mutant residues from the Psc3 hypomorphs near the interacting regions with Rad21 or Rec8, possibly ruling out any structural basis for the contrasting rescue phenotypes observed.

### Chromatin-bound Rec8-Psc3 mutant cohesin complex persists despite restrictive conditions to trigger Psc3 depletion

In order to test whether the rescue in viability was somehow due to restoration of centromeric cohesion, we determined the chromatin-bound occupancy of Psc3-4ts mutant protein and Rec8 in these cells at the permissive and non-permissive temperatures. Surprisingly, ChIP data showed that both Psc3-4ts and Rec8 were enriched to various degrees at the centromeric core, the flanking pericentromeric loci and the arm loci at both temperatures (Fig. 5A and Suppl. Fig. 3D). This suggests potential stabilization of the Rec8-Psc3-4ts cohesin complex, despite restrictive temperatures. Interestingly, upon *swi6Δ,* the enrichment of Psc3-4ts and Rec8 dropped overall but specifically at the pericentromeric loci, while being present at the centromeric core and the arms (Fig. 5B and Suppl. Fig. 3E).

**Figure 5.**
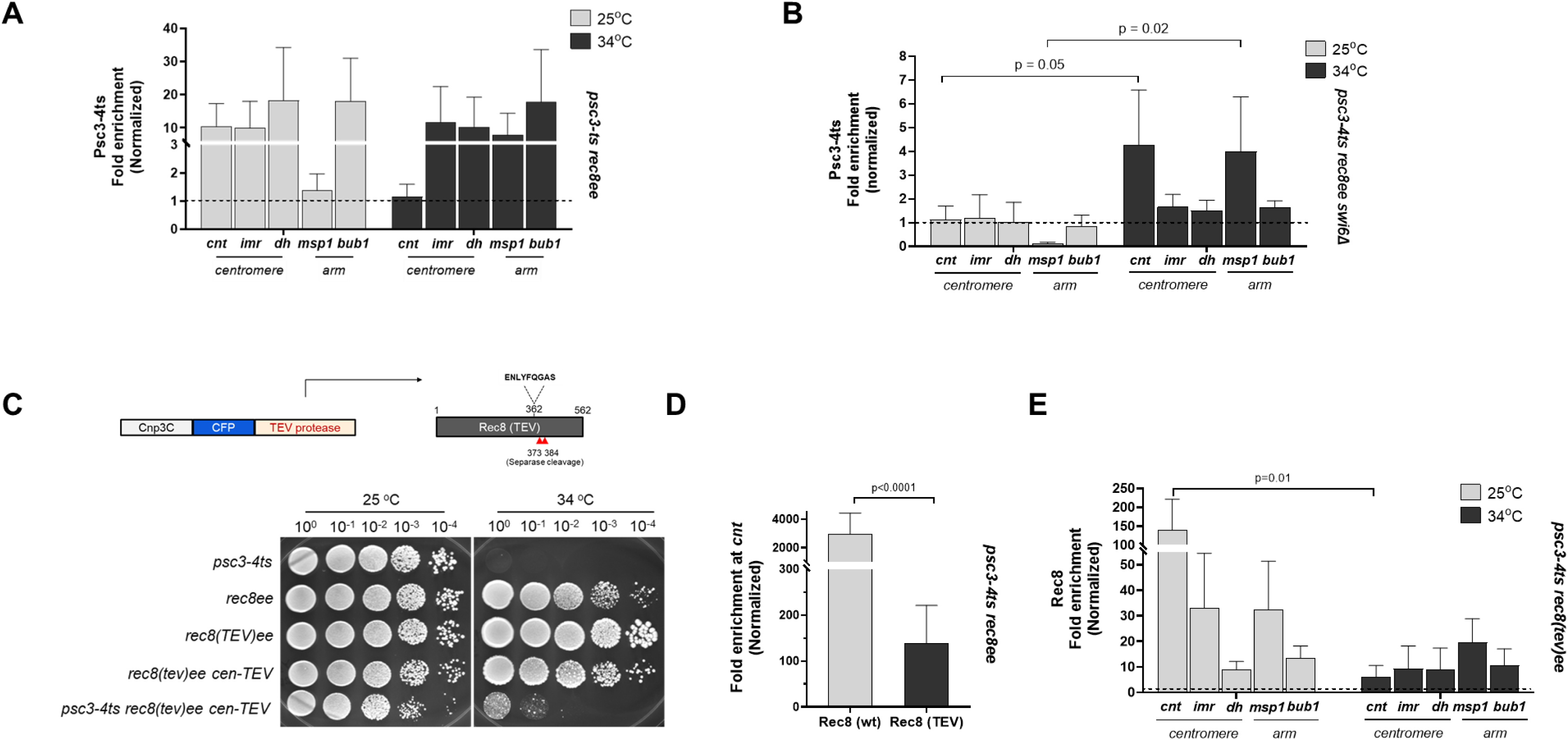
Two populations of Rec8-Psc3-ts complex exist at the centromeres similar to meiosis. A-B. ChIP qPCR for Psc3-4ts mutant in *psc3-4ts rec8ee* (A) and *psc3-4ts rec8ee swi6Δ* (B) at centromere and arm loci at 25 °C and 34 °C. Fold enrichment was calculated after normalizing with mitochondrial DNA locus as negative control. **C.** Spot assay for *psc3-4ts* in the presence of TEV protease (*cen-TEV*) cleavable Rec8 (*rec8(tev)ee* at 25 °C and 34 °C. A schematic for *rec8(tev)ee* and *cen-TEV* alleles is provided. **D.** Rec8 enrichment at the centromere core in *rec8(wt)ee* and *rec8(tev)ee* strains. **E.** ChIP qPCR for Rec8 in *psc3-4ts rec8(tev)ee cen-TEV* at centromere and arm loci at 25 °C and 34 °C. Fold enrichment was calculated after normalizing with mitochondrial DNA locus as negative control. **F.** Spot assay for *psc3-4ts* in the presence of TEV protease (*cen-TEV*) cleavable Rec8 (*rec8(tev)ee* with and without *swi6Δ* at 25 °C and 34 °C. For each experiment, atleast 3 biological repeats were performed. For panels A, B, D and E, bars represent the mean of at least three independent experiments performed in triplicates; error bars=SEM.

Since, Rec8-Psc3-4ts complex enriches at the centromere core, we tested if specifically removing the cohesins from this region could have an impact on cell survivability, similar to *swi6Δ* (Fig. 4A). We used a previously described system of a TEV protease cleavable allele of *rec8 (rec8(tev)ee)* and expressed the TEV protease as a fusion with the Cnp3C (*cen-TEV* allele) that localizes to the kinetochore at the centromere core^23^ (Fig. 5C). In the presence of *rec8(tev)ee,* the *psc3-4ts* cells still showed rescue of growth at 34 °C, however it was reduced, similar to that seen in *psc3-4ts rec8ee swi6Δ* (Fig. 5C, 4A). This clearly exhibits that the two populations of Rec8-Psc3 – one at the centromeric core and the other at the pericentromeric heterochromatin are individually sufficient towards cell survival, but the growth is largely compromised. ChIP analysis for Rec8 enrichment in the presence of the *rec8(tev)ee* allele at the centromeric loci, showed significant reduction in chromatin-bound levels of Rec8 at the centromere core (Fig. 5D). The Rec8 enrichment at the pericentromeric loci was still present, although reduced at 34 °C, supporting the partial rescue phenotype (Fig. 5E). These data confirm existence of two populations of Rec8-Psc3-4ts cohesin complexes at the centromeric core and the pericentromeric regions that can be formed independent of each other and support centromere cohesion.

### Rec8 can prolong stability of mutant Psc3 protein to form a functional cohesin complex

Chromatin enrichment of Psc3-4ts protein even at 34 °C, possibly suggests enhanced stability of a sub-fraction of Psc3-4ts mutant that remains in complex with Rec8, even at restrictive conditions that otherwise favor protein destabilization (Fig. 5A, 5B). To test this, we first determined the levels of Psc3 in the *psc3-4ts* mutant and found that it shows near wild-type protein levels, unlike the other two alleles, *psc3-2ts* and *psc3-3ts* (Suppl. Fig. 4A). Cells with just the *psc3-4ts* mutant allele, showed a near complete absence of the Psc3 protein when shifted to the restrictive 34 °C temperature compared to 25 °C (Fig. 6A). Interestingly, when *psc3-4ts rec8ee* cells, both in the presence and absence of Swi6, were grown at 34 °C, a significant amount of Psc3 protein was still present (Fig. 6A). The same persistence of mutant Psc3 protein was not observed, when Rad21 was over-expressed to similar levels as Rec8, supporting the previous observation of absence of rescue in cell viability (Suppl. Fig. 4B and Fig. 4C). This strongly suggested that the restoration of cell viability by Rec8 in the absence of Rec11 and conditional inactivation of Psc3, was brought about by a stabilized fraction of mutant Psc3 protein in these cells.

**Figure 6.**
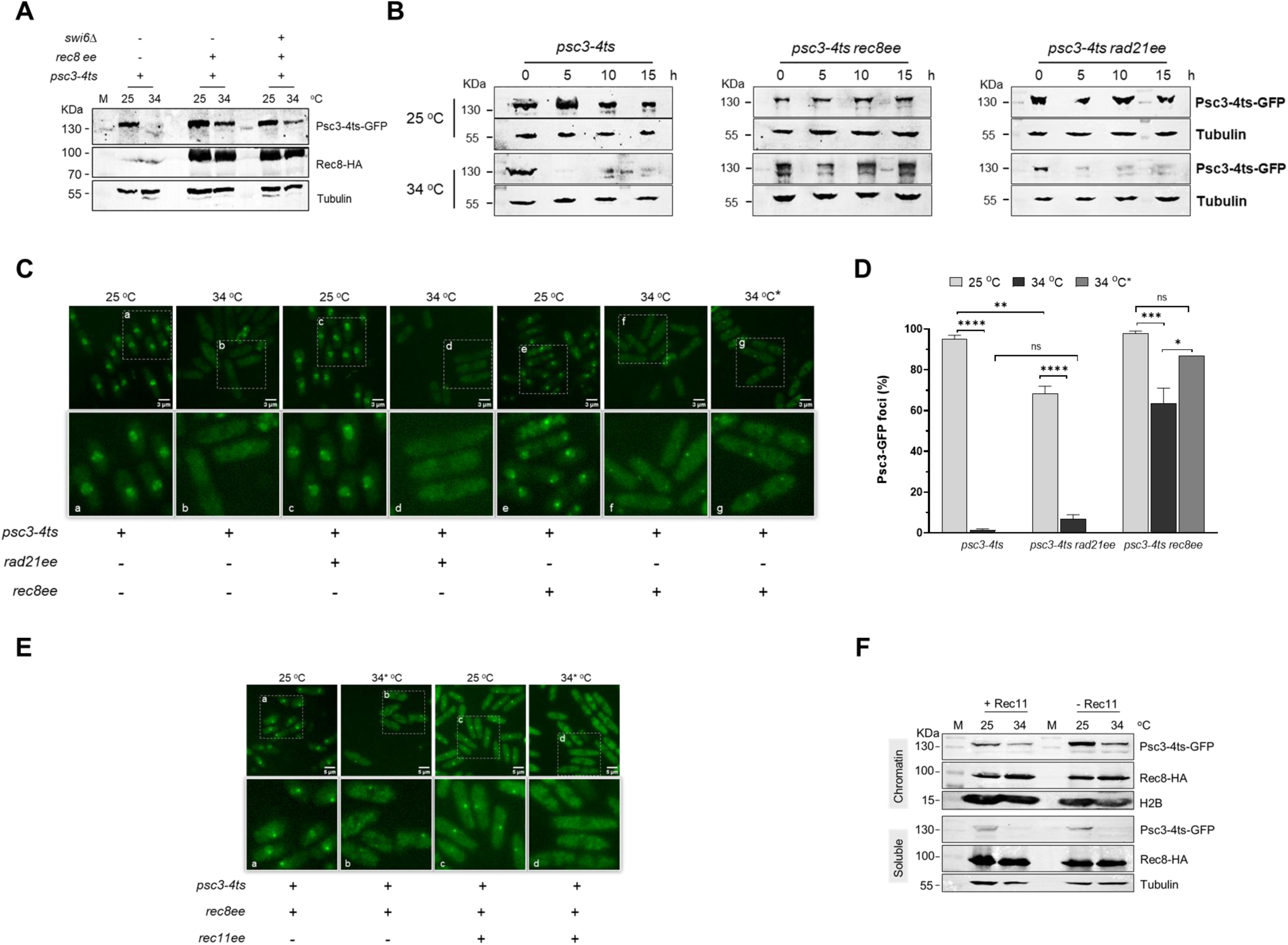
Psc3-ts mutant protein persists at restrictive temperature in the presence of Rec8 but not Rad21. **A.** Western blot to determine expression levels of Psc3-4ts mutant and Rec8 at 25 °C and 34 °C in the mentioned genotypes. The blots were probed with anti-GFP, anti-HA and anti-Tubulin antibodies. **B.** Time-course experiment to determine expression levels for Psc3-4ts mutant protein at 25 °C and 34 °C in non-cycling cells. The blots were probed with anti-GFP and anti-Tubulin antibodies. M is ladder. **C.** Representative images for Psc3-GFP foci formation for the *psc3-4ts* mutant in presence of *rad21ee* and *rec8ee* at 25 °C and 34 °C. 34 °C* represents cells grown at 34 °C completely, while the other cells were grown at 25 °C and shifted to 34 °C for 8 h. Scale: 3 µm. **D.** Graph quantifying the percentage of Psc3-GFP foci in cells analyzed in panel C. n=2, the error bars represent range and 200 cells were analyzed for each condition for each strain. Data analyzed by Two-way ANOVA test (Tukey’s multiple comparisons test) ****p <0.0001; ***p <0.001; ** p<0.01; *p <0.05; ns p >0.05. **E.** Representative images for Psc3-GFP foci formation for the *psc3-4ts rec8ee* with and without *rec11ee* at 25 °C and 34 °C. Scale: 5 µm. **F.** Expression levels of Psc3-4ts mutant and Rec8 at 25 °C and 34 °C in the presence and absence of Rec11. The chromatin and soluble fractions were analyzed separately. The blots were probed with anti-GFP, anti-HA, anti-H2B and anti-Tubulin antibodies. For each experiment, atleast 3 biological repeats were performed.

We further tested the stability of the mutant Psc3 in the presence of overexpressed Rec8 and Rad21 in non-cycling cells (starved at G1) to understand the protein turnover dynamics. As expected, the mutant Psc3 protein was stable across nearly all the time-points tested in *psc3-4ts* cells, independent of the overexpressed kleisin partner protein at the permissive temperature (Fig. 6B). However, when cells were grown at the restrictive temperature (34 °C), the levels of Psc3 became undetectable by 5h for both the *psc3-4ts* strains with normal and overexpressed Rad21 kleisin (Fig. 6B). However, Psc3 levels continued to persist till around 15h post-shift to 34 °C when Rec8 was expressed in these cells (Fig. 6B). Levels of overexpressed Rec8 and Rad21 were largely still present, although reduced across the time-course at both the temperatures (Suppl. Fig. 4C).

Furthermore, we also checked for the formation of Psc3-GFP foci in proliferating cells, which are attributed to the centromeric population due to centromere clustering, at both 25 °C and 34 °C in different *psc3-4ts* strains. As expected, Psc3-4ts-GFP foci was seen only at 25 °C and there was a complete absence of Psc3-4ts-GFP foci at 34 °C (Fig. 6C (a,b), 6D). When Rad21 was overexpressed, there was still a near absence of Psc3-4ts-GFP foci, similar to the reduction in protein levels (Fig. 6C (c, d), 6D, 6B). However, upon Rec8 expression, we saw comparatively better Psc3-4ts-GFP foci formation even at 25 °C and in agreement with the protein levels above, continued expression and formation of Psc3-4ts-GFP foci, when the cells were shifted to 34 °C for 8 h (Fig. 6C (e, f), 6D). Interestingly, we saw an increase in the fraction of cells with Psc3-4ts-GFP foci formation when the cells were grown at 34 °C continuously, possibly indicating a higher fraction of dividing cells in the culture with the stable Psc3-4ts sub-population, compared to when they are shifted to the restrictive temperature for only a few hours (Fig. 6C (e-g), 6D). These data support our inferences from above that the residual population of Psc3-4ts mutant protein must be a sub-population of Psc3, which is in a stable complex with Rec8 and deposited at the centromeres, both at the core and pericentromeric heterochromatin, as confirmed by ChIP (Fig. 5A, 5B).

We also tested whether this stabilization of mutant Psc3 by Rec8 could be maintained upon co-expression of Rec11 in *psc3-4ts* cells. We could still observe the Psc3-4ts-GFP foci at 25 °C even in the presence of Rec11, confirming a lack of interference, likely due to exclusion of Rec11 from the centromeres when Psc3 is present (Fig. 6E(c)). However, at 34 °C the cells did not show any Psc3-4ts-GFP foci, although there was still residual amount of chromatin-bound Psc3 protein present (Fig. 6E(d); 6F). Loss of Psc3 mutant protein in the soluble fraction at 34 °C, indicated general destabilization of the Psc3-ts protein that is not part of a chromatin-bound functional complex with Rec8 (Fig. 6F).

### The stabilization of mutant Psc3 by Rec8 can rescue segregation defects in meiosis but not in proliferating cells

We next assessed the effect of this stabilization of Psc3 mutants via Rec8 on segregation fidelity in both mitotically dividing cells and during meiosis. We chose *psc3-3ts* allele as it had the least amount of detectable Psc3 protein even at 25 °C (Suppl. Fig. 4A). As expected *psc3-3ts* mutant showed hypersensitivity to TBZ, which did not alter much upon further addition of *rec8ee* (Fig. 7A). Further removal of Swi6, increased the sensitivity to TBZ, which could be due to the reduced pericentromeric cohesin loading (Fig. 7A and Suppl. Fig. 3E). *rec8ee* on its own shows TBZ sensitivity, which could be due to its enrichment at the *cnt* core and dismantling of the kinetochore, as shown previously^25^. In order to test if reducing the Rec8 cohesins at the centromeric core could revert back the TBZ sensitivity, we used *rec8(TEV)ee* along with the Cnp3C-TEV protease (Suppl. Fig. 4D). However, unexpectedly, we observed slightly more sensitivity to TBZ, compared to *rec8ee* alone, indicating that mis-segregation in the presence of Rec8, could not just be attributed to the population of cohesins at the centromere core (Suppl. Fig. 4D).

**Figure 7.**
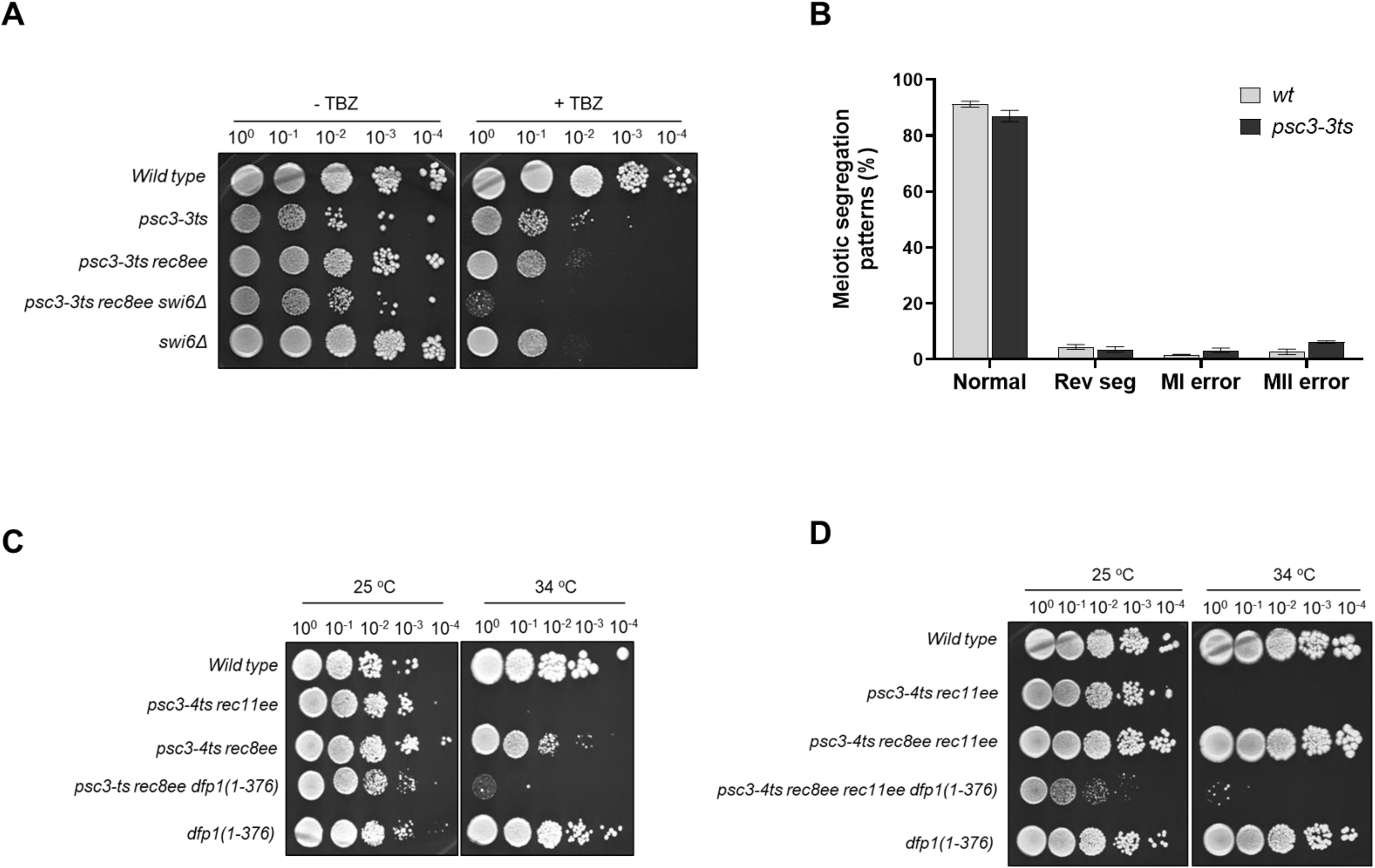
Rec8-Psc3-ts cohesin complexes at the centromere contribute to segregation defects and require DDK kinase for loading. **A.** Spot assay to assess hypersensitivity to Thiabendazole (TBZ) (0 and 10 μg/ml) in *psc3-3ts* and *psc3-3ts swi6Δ* backgrounds upon expression of Rec8. **B.** Meiotic segregation fidelity was determined via a fluorescence-based tetrad assay, described previously^30,31^. Graph showing the percentage of tetrads showing normal segregation, reverse segregation phenotype, meiosis I (MI) and meiosis II (MII) errors in wild type and *psc3-3ts* mutant. The error bars represent ±SEM. **C-D.** Spot assay for *psc3-4ts* in the presence of *rec8ee* (C) and *rec8ee rec11ee* (D) along with *dfp1(1-376)* truncation that affects recruitment of Hsk1-Dfp1 kinase complex to the centromeres at 25 °C and 34 °C. For each experiment, atleast 3 biological repeats were performed.

We tested the same *psc3-3ts* hypomorph for fidelity of meiotic segregation using a fluorescence-based assay described before^30,31^ (Suppl. Fig. 4E). *psc3Δ rec8ee rec11ee* cells show a high percentage of errors in meiosis II (MII) segregation^31^. We expected the *psc3-3ts* hypomorph to also show increased MII errors in meiosis, due to extremely low Psc3 protein levels, which would result in improper segregation of sister-chromatids at MII. However, we observed near normal meiotic segregation patterns in both MI and MII (Fig. 7B). Since, the Rec8-Rec11 complex in *psc3Δ rec8ee rec11ee* cells cannot rescue the segregation defects, we can infer that the *psc3-3ts* hypomorph must be getting stabilized in the presence of meiotically expressed Rec8, to show wild type chromosomal segregation, similar to what we have seen above in proliferating cells.

### Rec8-mediated cohesin loading at the centromere is dependent on DDK activity

Hsk1-Dfp1 kinase complex (DDK) is known to play a critical role in cohesin loading at the centromeres. In *S. pombe,* it is proposed to mediate the interaction between Swi6 and Psc3 and thereby help in cohesin loading at the pericentromeric heterochromatin^15,32^. In *S. cerevisiae*, the DDK mediates interaction of the cohesin loader with the inner kinetochore protein, to facilitate cohesin loading at the centromeres^17^. Therefore, we wondered if the Hsk1-Dfp1 kinase complex could contribute to the Rec8-mediated cohesin loading at both the centromeric core and pericentromeric heterochromatin. We tested a *dfp1(1-376)* C-terminal truncation mutant of Dfp1 that has been shown to abolish its interaction with Swi6 and results in reduced cohesin deposition and thereby chromosomal mis-segregation^15^. In the presence of *dfp1(1-376), psc3-4ts rec8ee* cells were nearly completely sensitive at 34 °C (Fig. 7C). This was also true for cells additionally expressing *rec11ee,* indicating that loading of both Rec8-Psc3-ts and Rec8-Rec11 complexes in Psc3 depletion conditions, was inhibited by loss of Hsk1-Dfp1 activity, at the centromeres (Fig. 7C, 7D). Although the *dfp1(1-376)* mutant lacks the PxVxL motif to interact with Swi6, it also lacks the C-terminal Zinc finger domain, which may play a role in its binding to other chromatin regions and consequently its broader effect on cohesin loading. Absence of DDK activity could explain loss of cohesin loading at the centromere core, if a mechanism similar to that in budding yeast exists in *S. pombe*.

## Discussion

Cohesin complexes play a fundamental role in maintaining genome stability by ensuring accurate chromosome segregation, yet the molecular rules that dictate whether all the cohesin complexes that assemble get loaded onto the chromatin, especially in the context of presence of multiple paralogs remain poorly understood. Here, we systematically explored the functional consequences of combining mitotic and meiotic cohesin subunits in proliferating *Schizosaccharomyces pombe* cells. Our results reveal that the composition of cohesin complexes strongly influences their ability to get deposited at the centromeres, the chromatin residence time, and ultimately their capacity to support faithful chromosome segregation as well as other critical processes such as gene regulation and genome organization. Importantly, we identify an unexpected flexibility in cohesin complex assembly and chromatin loading mediated by the meiotic kleisin Rec8, which allows cells to survive conditions where canonical cohesin loading factors are compromised.

### Differential functional plasticity provided by the two kleisin paralogs

One of the key observations of our study was that the hybrid cohesin complex composed of the mitotic kleisin Rad21 and the meiotic HAWK Rec11 was largely non-functional in proliferating cells that were depleted of the canonical HAWK partner Psc3. Defective centromeric localization rather than complex formation itself was the cause for cell inviability highlighting that all cohesin complexes assembled need not get efficiently loaded at all the critical chromosomal regions (Fig. 8). Another possible explanation could be the lower dwell times on the chromatin as seen for Rec11 in the absence of Rec8, which could prevent stable persistent binding. On the contrary, the meiotic kleisin Rec8 displayed remarkable flexibility in forming functional cohesin complexes irrespective of the HAWK partner, an intrinsic property derived from its role during meiosis, where multiple cohesin complexes must coexist and function in distinct chromosomal domains (Fig. 8). Another striking observation was the ability of Rec8 to stabilize chromatin-bound hypomorphic Psc3 mutants and restore their functionality, even under restrictive conditions that normally trigger mutant Psc3 degradation. The residence time of Rec8-Psc3 cohesins was observed to be long, especially at the heterochromatin, which could potentially contribute to the retention of Psc3 mutants for a longer time on the chromosomes and preventing faster degradation. We also established that two distinct populations of the Rec8-Psc3 mutant complexes—one at the centromeric core and another at pericentromeric heterochromatin—can independently contribute to cell survival. Selective removal of either population reduced growth but did not eliminate viability, indicating synergistic effects of the two cohesin pools in providing stability.

**Figure 8.**
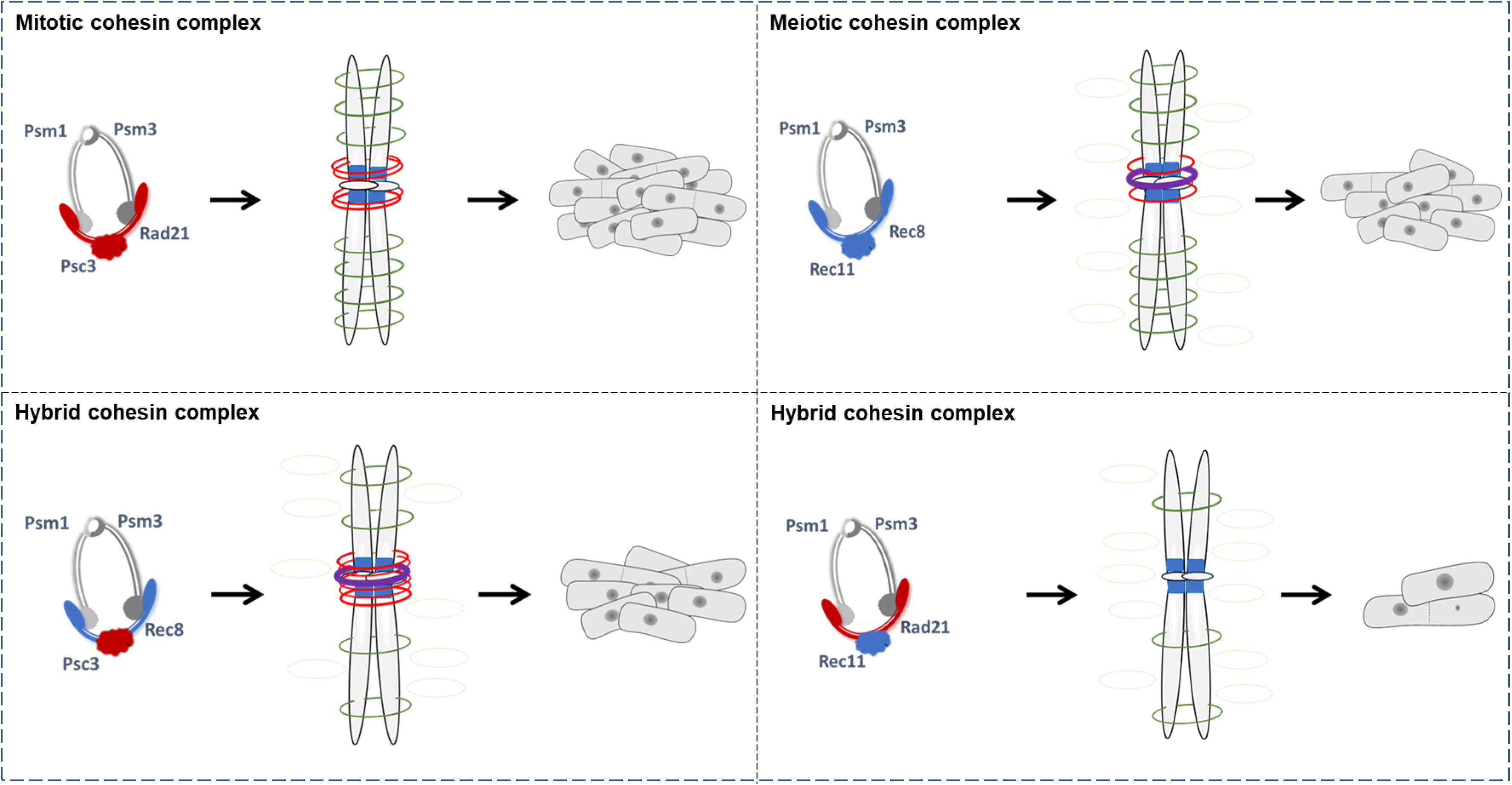
Model summarizing the kleisin partner preferences and cohesin loading patterns that dictate cellular viability. The four possible cohesin complexes that can be formed in *S. pombe* utilizing all the paralogs are shown. The mitotic Rad21-Psc3 is the canonical complex supporting complete growth and high fidelity of segregation. The completely meiotic Rec8-Rec11 complex shows better loading in the chromosomal arms and minimal pericentromeric loading, leading to defects in chromosomal segregation. The two hybrid complexes, Rad21-Rec11 and Rec8-Psc3 show opposite phenotypes. Insufficient cohesin loading in the presence of Rad21-Rec11 leads to cell death, whereas Rec8-Psc3 shows robust centromeric loading but leads to segregation defects and hence slightly compromised growth. The purple cohesin ring represents the Rec8 cohesin loaded at the centromere core regions. The red cohesin ring is the population loaded at the pericentromeric heterochromatin, which is marked in blue.

### Cohesin composition influences chromosomal segregation fidelity

Although alternative cohesin complexes could support cell survival in otherwise non-permissive conditions, they did not fully recapitulate the fidelity of canonical Rad21–Psc3 cohesins during chromosomal segregation (Fig. 8). Cells relying on meiotic cohesin complexes (Rec8-Psc3 and/or Rec8-Rec11) for survival exhibited increased sensitivity to the spindle poison Thiabendazole, thereby indicating higher rates of chromosome mis-segregation. Deposition of Rec8 complexes at the centromere core has been shown previously to increase kinetochore instability and generate uniparental disomy that may contribute to genomic instability and potentially tumorigenesis^21,25^. Although removal of just the centromere core population of Rec8 did not restore segregation to wild type levels in our study, detrimental consequences of Rec8 cohesin complexes on other cellular processes, cannot be ruled out. We also find significantly longer residence times on heterochromatin compared to euchromatin for Rec8-Psc3, which is unlike Rec8–Rec11 complexes that exhibited shorter residence times at heterochromatin, in the absence of Psc3. Reduced stability of Rec8–Rec11 complexes at the pericentromeric heterochromatin may lead to weaker centromeric cohesion, which could elevate chromosome segregation errors in cells that completely rely on these complexes, in the absence of the canonical cohesins (Fig. 8). Hence, the type of cohesin loaded across the chromatin does have a strong consequence on chromatin-binding kinetics and thereby the strength of cohesion at specific genomic regions that atleast affects chromosomal segregation as tested in this study and may disrupt genome architecture and transcriptional regulation.

### Implications for cohesin dysregulation in cancer

Human cancers express a class of proteins known as the Cancer/Testis Antigens (CTA) that typically are expressed only in the germ cells and meiosis-specific cohesin subunits are a consistent part of this group^10,33–35^. Aberrant expression of meiotic paralogs such as REC8, STAG3, and SMC1β has been documented in multiple cancer types, where the coexistence of mitotic and meiotic cohesin subunits is inevitable^34^. Our work provides a conceptual framework for understanding the potential consequences of such dysregulation. We show that cells can assemble a variety of hybrid cohesin complexes when multiple paralogs are present, some of which can compensate for loss of canonical cohesin components but at a fitness cost. Moreover, cohesin subunits such as STAG2 (Psc3 ortholog) are frequently mutated in cancers and their functional inactivation can reduce sufficient cohesin occupancy on chromosomes that can have consequences on cellular growth^36–39^. However, there are reports that find that inactivating mutations in STAG2 do not always result in loss of cohesin functions and compensatory mechanisms have been suggested to play a role^40–42^. We propose that the flexibility provided by meiotic kleisins such as REC8 may allow cancer cells harboring such cohesin mutations to maintain minimal cohesion and survive despite compromised canonical pathways. However, these alternative complexes may also introduce chromosome segregation errors due to altered chromatin binding dynamics, potentially contributing to genomic instability and aneuploidy that are frequently observed in tumors^36,43,44^. Therefore, the formation of hybrid cohesin complexes may represent both a survival strategy and a source of chromosomal instability in human cancers.

Taken together, our findings suggest a model in which cohesin complex functionality is determined not only by the availability of individual subunits but also by intrinsic compatibility between kleisin and HAWK partners. While canonical Rad21–Psc3 cohesins provide optimal centromeric cohesion and segregation fidelity in proliferating cells, the meiotic kleisin Rec8 provides flexibility that allows formation of alternative complexes capable of sustaining cell survival when canonical loading pathways fail, although at significant fitness costs (Fig. 8). This work therefore uncovers previously unappreciated rules governing cohesin complex assembly and chromatin loading. Understanding these principles are critical for elucidating how cohesin diversity contributes to genome organization in normal development and how its dysregulation promotes genome instability in disease.

## Material and Methods

### Strains and genetic methods

The genotypes of the *S. pombe* strains generated and used in this study are described in Supplementary Table 1. The *S. pombe* strains were generated using standard genetic crosses and random spore analysis^45^. Strains were grown at 30 ⁰C or 25 ⁰C (for temperature sensitive-ts strains), and cultured mainly in YEA rich medium or supplemented minimal (EMM2) medium. Oligonucleotides and plasmids used in the study are described in Supplementary Tables 2 and 3, respectively. To generate the ectopically expressing Rec11 and Rec11-CD strains, *ura4+-Padh1* construct was amplified from plasmid ME27 (FYP4963 obtained from NBRP, Japan) with primers MO53 and MO54 replacing the native promoter of *rec11* via lithium acetate transformation method, as described previously^46^. *rec8-HaloTag* and *rec11-HaloTag* alleles were generated by introducing HaloTag-HygR and linker-HaloTag-HygR at the 3’-UTR of *rec8* and *rec11* genes, respectively. The amplicons were generated via overlap PCR using primers MO341-344 (for Rec8-HaloTag) and MO468, MO375, MO471 and MO377 (for Rec11-HaloTag), respectively, as described previously^47^. Colony PCRs were performed for genotyping.

### Spot Assay

Strains were freshly streaked on YES (Yeast Extract Supplemented) agar plates and single colonies were mixed in 100 μl freshly autoclaved distilled water. The OD_600_ of all the strains were maintained between 0.75-0.85. The cell suspensions were serially diluted 10-fold till 10^-4^ dilution and 7-10 µl of each dilution was spotted on YES agar plates and incubated at 25 °C and 34 °C for restrictive temperature growth assay. For Thiabendazole sensitivity assays, the strains were spotted on YES agar plates containing 0, 7.5 or 10 μg/ml TBZ drug and incubated at 25 °C.

### Growth Assay

A single colony was inoculated in 5 mL of YES liquid medium. The cultures were allowed to grow at 25 °C till saturation. The cells were sub-cultured in 3 ml YES until they reached an OD_600_ of 0.6-0.7 (log phase). The secondary culture was then diluted to a starting OD_600_ of 0.03-0.05 and incubated at 25 °C. Absorbance was measured every 6 h until saturation was reached.

### Western blot analysis

For checking protein expression, strains were cultured in YES media and harvested at OD_600_ – 0.7-0.8. To check for levels of Psc3-ts proteins, the respective strains were cultured at 25 ⁰C till OD_600_ – 0.3-0.4 and then shifted to 34 ⁰C for 8-10hrs. To check for the turnover of Psc3-4ts protein at 34 ⁰C, across various time-points, the strains were cultured in Edinburgh Minimal Media 2 (EMM2) containing NH4Cl as nitrogen source. Cultures were grown at 25 ⁰C till OD_600_ – 0.3-0.4 and then shifted to a nitrogen deficient EMM2 media. The cultures were split and incubated at both the permissive (25 ⁰C) and non-permissive (34 ⁰C) temperatures. Cells were harvested at 0, 5, 10 and 15 h time points. Harvested cells were washed and lysed using 1.85M NaOH and β-mercaptoethanol. Proteins were precipitated using 50% Trichloroacetic acid (TCA) and resuspended using 1X Laemmeli buffer and neutralised by Tris-HCL (pH-8). Equal amounts of protein were loaded on a 7.5% SDS-PAGE gel and subsequently transferred to a PVDF membrane (Merck IPVH00010-IN). Blots were blocked using 5% skim milk solution and probed using 1:1000 dilutions of anti-GFP (Invitrogen MA515256), anti-HA (Invitrogen 26183), anti-FLAG (Sigma F1804), anti-HaloTag (Promega G9211), anti-tubulin (Invitrogen 62204) and 1:2000 dilutions of anti-Swi6 (BioAcademia Inc 63-101) and anti-Rad21 (BioAcademia Inc 63-139). Goat anti-mouse IgG H+L (Jackson Immunoresearch 115-035-003), Goat anti-rabbit IgG H+L (Jackson Immunoresearch 111-035-003), Mouse anti-rabbit Light chain specific IgG (Jackson Immunoresearch 211-032-171) and Rabbit anti-mouse Light chain specific IgG (CST 58802S) were used as secondary antibodies and the blots were developed using Chemiluminescent substrate (Invitrogen 34577) and subsequent imaging were done on iBright CL1500 imaging system.

### Co-immunoprecipitation

Strains were grown in 50 ml YES media till log phase (OD_600_ - 0.7 - 0.8) and crosslinked using 0.1% formaldehyde solution [37% formaldehyde solution (Qualigens Q12755), 5mM HEPES-KOH, 550mM NaCl and 0.1mM EDTA] for 50 min at 30 ⁰C and then quenched by adding 250mM glycine for 10 min. Cells were harvested and washed once with autoclaved distilled water, follwed by 1X TBS and complete lysis buffer [50mM Tris-Cl pH-7.5, 150mM NaCl, 5mM MgCl2, 5% glycerol, 1% NP40, freshly added 1X protease inhibitor cocktail (Sigma 4693159001), 60mM ꞵ-glycerophosphate (Sigma G5422), 1mM DTT and 2mM PMSF]. Cells were then resuspended in 400 μl of complete lysis buffer and lysed in a mortar pestle under liquid N2 till 70-80% lysing efficiency was observed under the microscope. The lysed cells were collected and spun down at 14000 rpm for 30 min at 4 ⁰C. The supernatant, which is the soluble fraction, was collected. The pellet was washed thrice with autoclaved distilled water to remove any soluble proteins. The pellet was then resuspended in 250 μl of complete lysis buffer and sonicated using Diagenode Bioruptor® Plus (15 cycles, 30 sec on/off, high amp) to shear the chromatin and release the chromatin-bound proteins. The sonicated fraction was then spun down at 14000 rpm for 10 min at 4 ⁰C, and the chromatin fraction was collected. For immunoprecipitation (IP), 5 μg of anti-GFP, anti-HA or anti-FLAG antibodies, 6.25 μg of anti-HaloTag antibody or 2.5 μl of anti-Swi6 or anti-Rad21 sera were used. 20 μl protein G-coated Dynabeads (Invitrogen 10003D) were incubated with the required antibodies overnight at 4 ⁰C. Antibody-coated beads were then incubated with the lysates for 3 h at 4 ⁰C. The beads were then washed thrice with lysis buffer backbone for 5 min each and eluted in 30 μl 2X Laemmeli buffer at 99 ⁰C for 5 min. The IP eluate and 2-5% input was loaded on a 7.5% SDS-PAGE gel and western blotting was done for subsequent analysis.

### Chromatin Immunoprecipitation

200 ml cultures growing in log phase (OD_600_- 0.7 - 0.8) were harvested and crosslinked using 1.1% formaldehyde solution [16% formaldehyde, methanol free (Invitrogen 28908)], 5mM HEPES-KOH, 550mM NaCl and 0.1mM EDTA for 50 min at room temperature. Excess formaldehyde was quenched using 250mM glycine for 10 min. Cells were washed twice with 1X TBS and then resuspended in 500 μl of Breaking buffer (100mM Tris-Cl pH-8, 20% glycerol and freshly added 1mM PMSF). Cells were lysed with mortar pestle under liquid Nitorgen till 70-80% lysing efficiency observed under microscope. Lysed cells were collected and washed 4 times with FA buffer (50mM HEPES-KOH, 150mM NaCl, 1mM EDTA, 1% Triton X-100, 0.1% Sodium deoxycholate and freshly added 1X Protease inhibitor cocktail). Pellets were resuspended in 300 μul FA buffer and sonicated using Diagenode Bioruptor® Plus (25 cycles, 30 sec on/off, high amp) till 200-700 bp chromatin shearing efficiency was achieved. 1.2ml FA buffer was added to the sonicated cells and finally spun down at 1500 rcf for 10 min at 4 ⁰C and a clear supernatant was collected. 10% of the lysate was kept aside and the remaining chromatin was first precleared using protein G-coated Dynabeads and then immunoprecipitated using the appropriate antibody overnight at 4 ⁰C with rotation. Antibody-conjugated lysates were then finally incubated with beads for 4 hr at 4 ⁰C. Beads were collected and then washed twice for 5 min each with FA buffer (50mM HEPES-KOH, 150mM NaCl, 1mM EDTA, 1% Triton X-100, 0.1% Sodium deoxycholate), FA-HS buffer (50mM HEPES-KOH, 500mM NaCl, 1mM EDTA, 1% Triton X-100, 0.1% Sodium deoxycholate), RIPA buffer (10mM Tris pH-8, 0.25M LiCl, 0.5% NP40, 0.5% Sodium deoxycholate and 1mM EDTA) and TE buffer (10mM Tris pH-8 and 1mM EDTA). Elution was done twice with 2X Stop Buffer (20mM Tris pH-8, 100mM NaCl, 20mM EDTA and 1%SDS) at 65 ⁰C for 30min. The eluted sample as well as 10% input were de-crosslinked at 65 ⁰C overnight and treated sequentially with RNase A (10 μg) (HiMedia DS0003) and proteinase K (20 μg) (Invitrogen 25530015) for 2 h at 37 ⁰C. DNA was then purified using Qiagen QIAquick PCR purification columns (Qiagen 28115). TB Green Premix Ex Taq II buffer mix (Takara RR820A) was used to set up quantitative PCR reactions for the purified input and immunoprecipitated DNA and analysed. Mouse (Merck 12-371) and Rabbit (Merck 12-370) normal IgGs were used as controls to check for background pulldown for all the antibodies in each experiment. Fold enrichment was calculated by subtracting the IgG values and a negative control region (mitochondrial loci - MitoNegative) to get the normalized value for each tested locus (2^-ddCt^).

### AlphaFold analysis

Amino acid sequences for Rec8, Rad21, Psc3 and Rec11 were taken from PomBAase^48^ and Alphafold 3 was used to predict their structures^49^. Amino acid residues 284-386 and 356-439 for Rec8 and Rad21, respectively, were separately tested for interactions with Psc3 and Rec11. Higher iPTM values were used as an indicator for higher confidence in interaction probability.

### Single-molecule imaging

The cells were streaked on YESA plate from the glycerol stock and incubated at 30 ℃ for 72 hrs. A single colony was inoculated in 5 ml EMM media (Edinburgh’s Minimal Media, MP Biomedicals, Cat no. 4110-012) with required supplements and grown at 30 ℃ under shaking condition (230 rpm) for 48 hr. 50 μl of this culture was inoculated in 3ml fresh EMM with required supplements and grown for 8-10 hr (until log phase) at 30°C under shaking conditions (230 rpm). 1 ml culture was harvested and JF646-HaloTag Ligand (JF646-HTL, Promega, Cat no. GA112A) was added at 3-5 nM concentration for labelling Rec11-HaloTag and at 20-25 nM concentration for labelling Rec8-HaloTag. The cultures were kept for shaking for 30 min at 30℃. Cells were pelleted down by centrifugation (3000 RPM for 2 min) and washed twice with 1 ml of fresh EMM media with supplements to remove unbound JF646-HTL. Cells were finally resuspended in 20 μl of EMM media with supplements and 3 μl of this suspension was placed on cover glass with thickness 1.5μm (corning, cat no.: 2980-246, size 24×60mm), covered by an EMM agarose pad (EMM media with supplements +2% Seakem Agarose (Lonza, Cat no. 50004), size: 8×8mm)), and imaged under a Leica Dmi8 infinity TIRF microscope using Highly Inclined and Laminated Optical Sheet (HILO) illumination. From each agarose pad, cells were imaged for 60 min.

Single-molecule imaging was performed using a Leica (Model: Dmi8 automated) infinity TIRF microscope, equipped with a 100X/1.47 NA oil immersion TIRF objective lens, a Photometrics Prime 95B sCMOS camera and a 150mW 638nm laser for HILO illumination, controlled by Leica LAS X software version 3.10.0. Cells were focused using the FITC channel under the wide-field illumination, and the best field was selected based on the fluorescence from Swi6-GFP channel. The time-lapse movies were acquired (in a single focal plane of 256×256 pixels) with 50ms exposure, 150ms time interval for 200 frames with 30% laser power (0.1mW at the objective under HILO illumination)^26,47^.

### Single-molecule tracking and data analysis

Single-molecule tracking was performed using the MATLAB-based MatlabTrack software package as described previously^26,50^. Raw image stacks (.tif files) were imported into MATLAB and processed using the automated analysis pipeline implemented in the software. Movies were subjected to band-pass filtering (1–5 pixels) to reduce background noise and enhance diffraction-limited spots. Single-molecule spots were identified by intensity thresholding, followed by detection of local maxima (threshold: 2.5–4.5; window size: 7 pixels) and two-dimensional Gaussian fitting to determine particle positions.

Particles were linked into trajectories using a nearest-neighbor tracking algorithm, allowing a maximum frame-to-frame displacement of 6 pixels. Only trajectories of at least 4 consecutive frames were retained, and gaps of up to 4 frames were closed to account for transient fluorophore blinking. Bound molecules were defined using parameters previously established for chromatin-bound histone Hht1-HaloTag in *Schizosaccharomyces pombe* (R_min = 0.54 µm, R_max = 0.81 µm, N_min = 8 frames)^47^.

Survival probabilities were calculated as 1 – CDF of dwell times and fitted with a double-exponential decay model to estimate residence times and the fractions of slow- and fast-moving populations, corresponding to specifically and non-specifically bound molecules, respectively. Photobleaching correction was applied by normalizing survival distributions to the bleaching curve obtained from the decay in the number of detected molecules over time, as described previously^50^. Swi6–GFP foci represents heterochromatin, so the regions of interests (ROIs) were drawn around these foci to track the Rec8 and Rec11 single-molecules over the heterochromatin, whereas, the nuclear ROIs excluding these Swi6-GFP foci were made to track the Rec8 and Rec11 over the euchromatin.

### Fluorescence microscopy and live cell imaging

For determining the Psc3-4ts-GFP foci, cells were streaked on YESA plate from the glycerol stock and incubated at 25°C for 4-5 days. For primary culture, a single colony was grown at 25°C for 48 hours in 5 ml EMM with required supplements under shaking condition (230 rpm). Saturated cultures were diluted into secondary cultures to an initial optical density (OD₆₀₀) of ∼0.05–0.1 and incubated at 25°C until mid-log phase (OD₆₀₀ ∼0.3–0.4). Cultures were then split into two equal portions: one half was maintained at 25°C, while the other was shifted to 34 °C and incubated for an additional 8 h. In parallel, an additional secondary culture was prepared from the primary culture and directly incubated at 34°C until OD₆₀₀ reaches to ∼0.8.

Following incubation, cells were imaged live under EMM agarose pads to assess Psc3–GFP foci formation.

All fluorescence images were acquired using a Leica DMi8 inverted fluorescence microscope equipped with a 63× oil-immersion objective (NA = 1.40), LED illumination (475 nm for GFP), and a 16-bit Prime 95B sCMOS camera (Teledyne Photometrics), controlled by Leica LAS X software. Z-stack images were acquired according to Nyquist sampling criteria. Images were processed using ImageJ for brightness and contrast adjustment, maximum-intensity projection, and cell counting. Identical acquisition settings were used for all experimental conditions.

For assessing the meiotic segregation fidelity, the parental strains were tagged with different fluorophores, tdTomato and mCerulean, expressed under a spore autonomous promotor near *Cen1*. The cells were then spotted on SPA sporulating media at 25 °C for 2-3 days for inducing meiosis. The mating mix was then collected and asci were imaged using LeicaDM6 upright fluorescence microscope equipped with a 63X oil-immersion objective using Texas Red and SBL filter. The experiment was repeated three times with each repeat containing atleast 100 tetrads. The images were analysed using Las X software.

## Acknowledgement

We thank Gopika Menon, Nikhil Venkatesh and Sarthak Ghosh for their technical help. We thank Dr. Girish Deshpande and MN lab members for their inputs on the manuscript and helpful discussions throughout the course of this study. We thank Dr. Gayathri Pananghat and Dr. Krishanpal Karmodiya for helpful discussions. We acknowledge Prof. Gerald R. Smith for sharing *S. pombe* strains. SB acknowledges fellowship from CSIR [09/0936(15178)/2022-EMR-I]. Purva acknowledges PMRF fellowship (0702610). OV is supported by University Grants Commission fellowship (221610144761). SS acknowledges fellowship from IISER Pune. AK is supported by UGC fellowship (191620186730). This study was supported by the DBT/Wellcome Trust India Alliance Intermediate Fellowship [Grant no. IA/I/23/1/506752] and Science and Engineering Research Board (SERB) - Promoting Opportunities for Women in Exploratory Research (POWER) Research grant [SPG/2022/000881] to MN. GM acknowledges funding from the Department of Biotechnology-Emerging Frontiers in Biotechnology (BT/PR54475/BSA/33/357/2024(CN21105). We also thank IISER Pune shared facilities including the Microscopy facility for infrastructural support. We thank the DBT-SAHAJ facility for Single-Molecule and Super-Resolution Imaging at IIT Hyderabad (BT/INF/22/SP53103/2024) for single-molecule imaging experiments.

## Author Contributions

M.N. conceived the study. M.N., S.B., and Purva designed the study and analyzed the data. S.B., Purva, A.D., A.K. S.S. and O.V. conducted the experiments. A.K. and G.M. analyzed the single molecule imaging data. M.N. and G.M. acquired resources and funding. M.N. wrote the original draft of the manuscript with inputs from other authors.

