## Supplementary Data for "Meiotic cohesin paralogs govern cell survival by exhibiting flexibility in partner choice"

**A**

| Mutant *psc3* allele | Mutations |
|---|---|
| *psc3-4ts* | C740R |
| *psc3-2ts* | K418R, I532T, S954P |
| *psc3-3ts* | L629P, T677A |

**B**

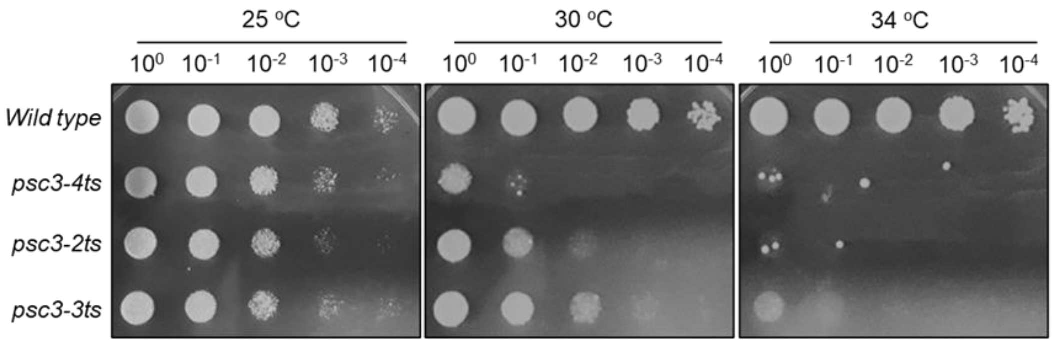


**C**

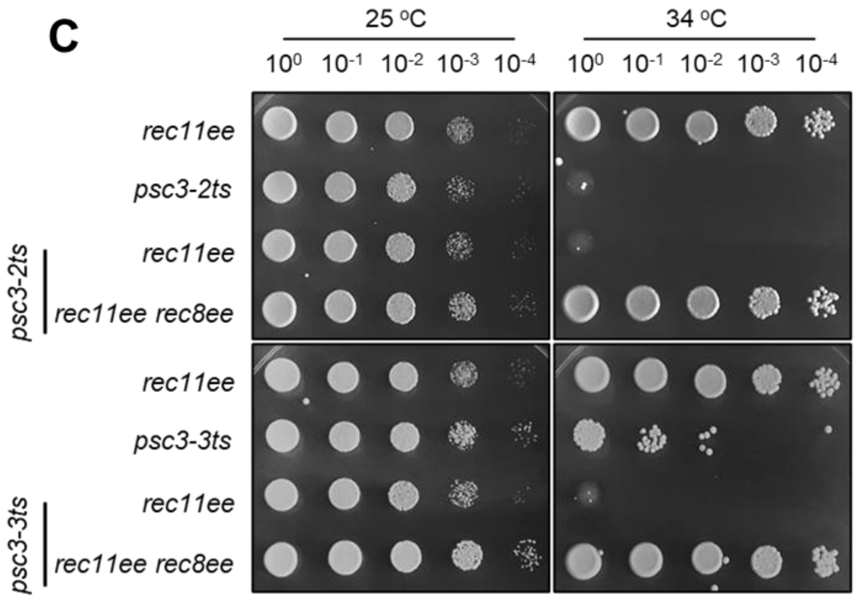


**D**

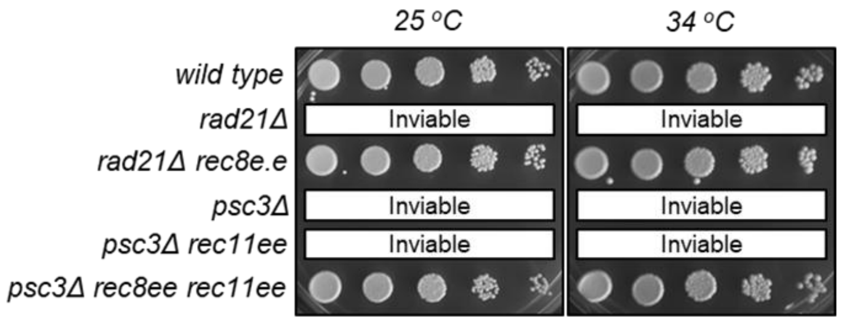


**E**

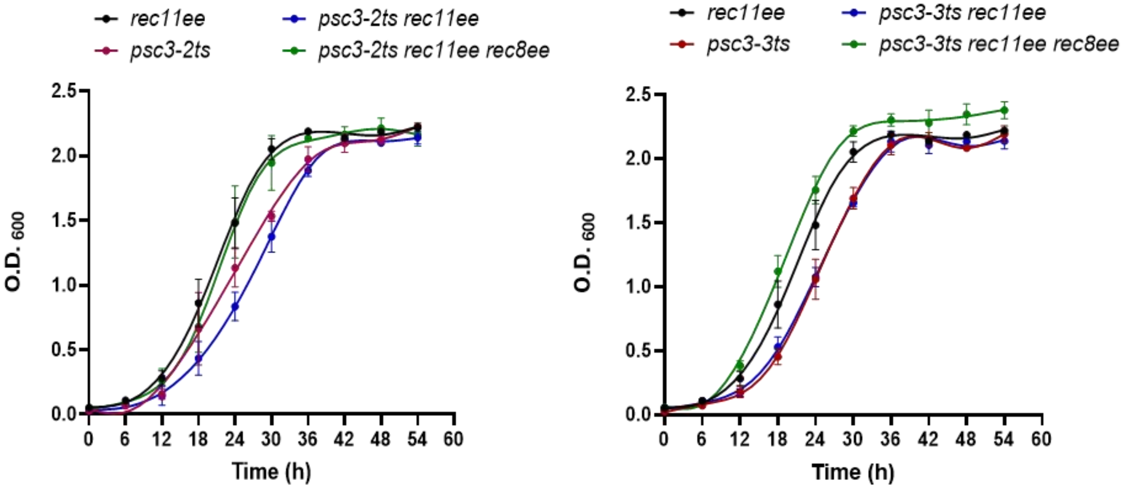


**F**

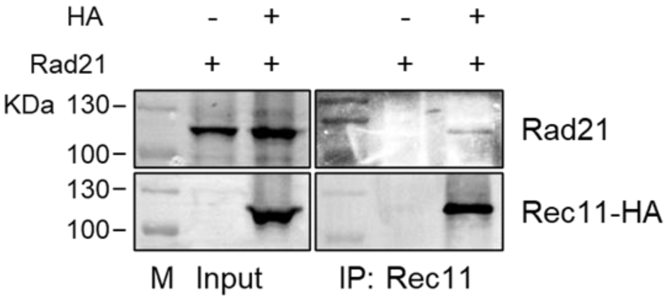


Suppl. Fig. 1

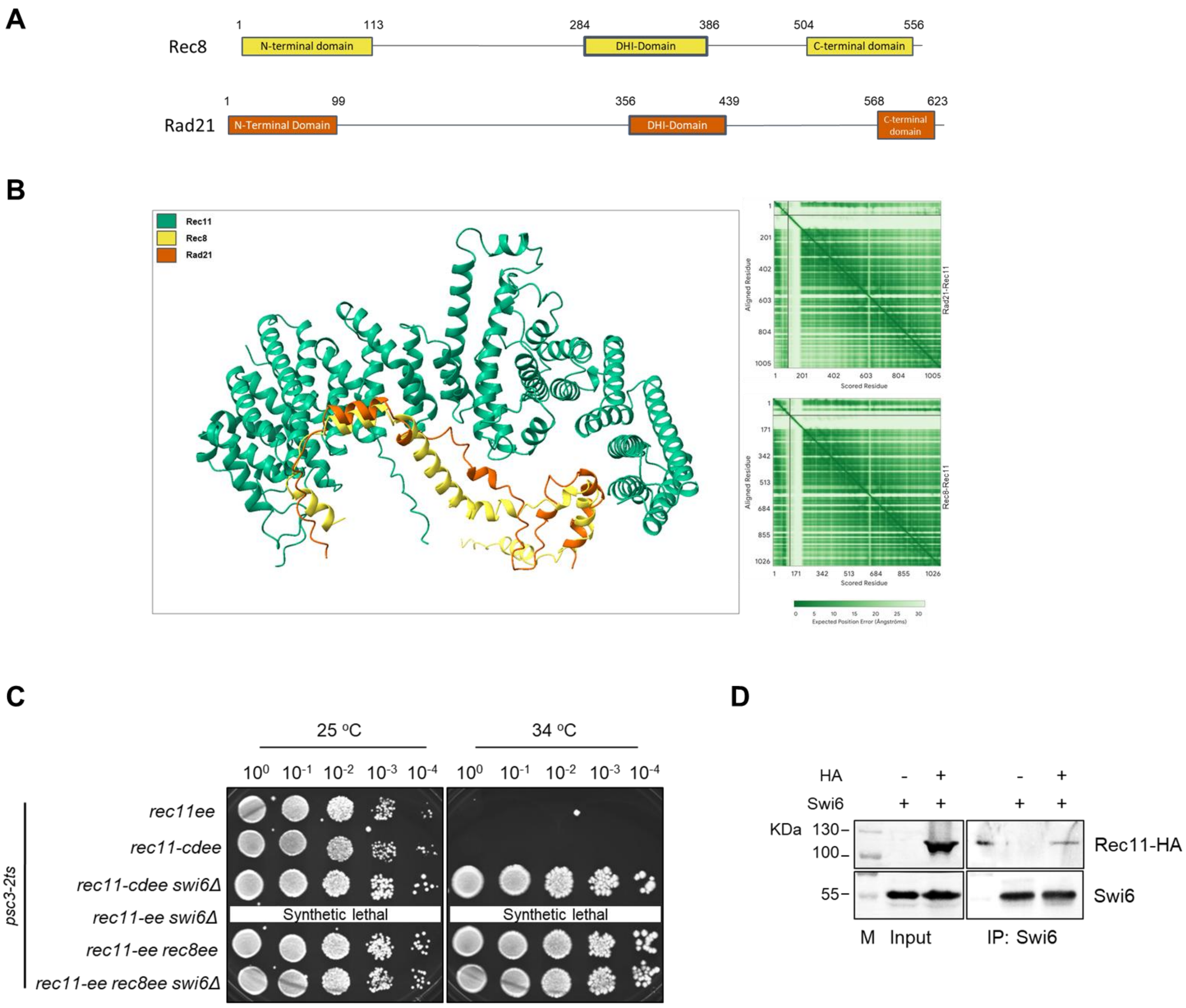
A
Rec8
1
113
284
386
504
556
N-terminal domain
DHI-Domain
C-terminal domain
Rad21
1
99
356
439
568
623
N-Terminal Domain
DHI-Domain
C-terminal domain
B
Rec11
Rec8
Rad21
Aligned Residue
Scored Residue
Rad21-Rec11
Rec8-Rec11
Expected Position Error (Ångströms)
C
25 °C
34 °C
$10^0$ $10^{-1}$ $10^{-2}$ $10^{-3}$ $10^{-4}$
psc3-2ts
rec11ee
rec11-cdee
rec11-cdee swi6Δ
rec11-ee swi6Δ
rec11-ee rec8ee
rec11-ee rec8ee swi6Δ
Synthetic lethal
D
HA
Swi6
KDa
130
100
55
Rec11-HA
Swi6
M
Input
IP: Swi6

Suppl. Fig. 2

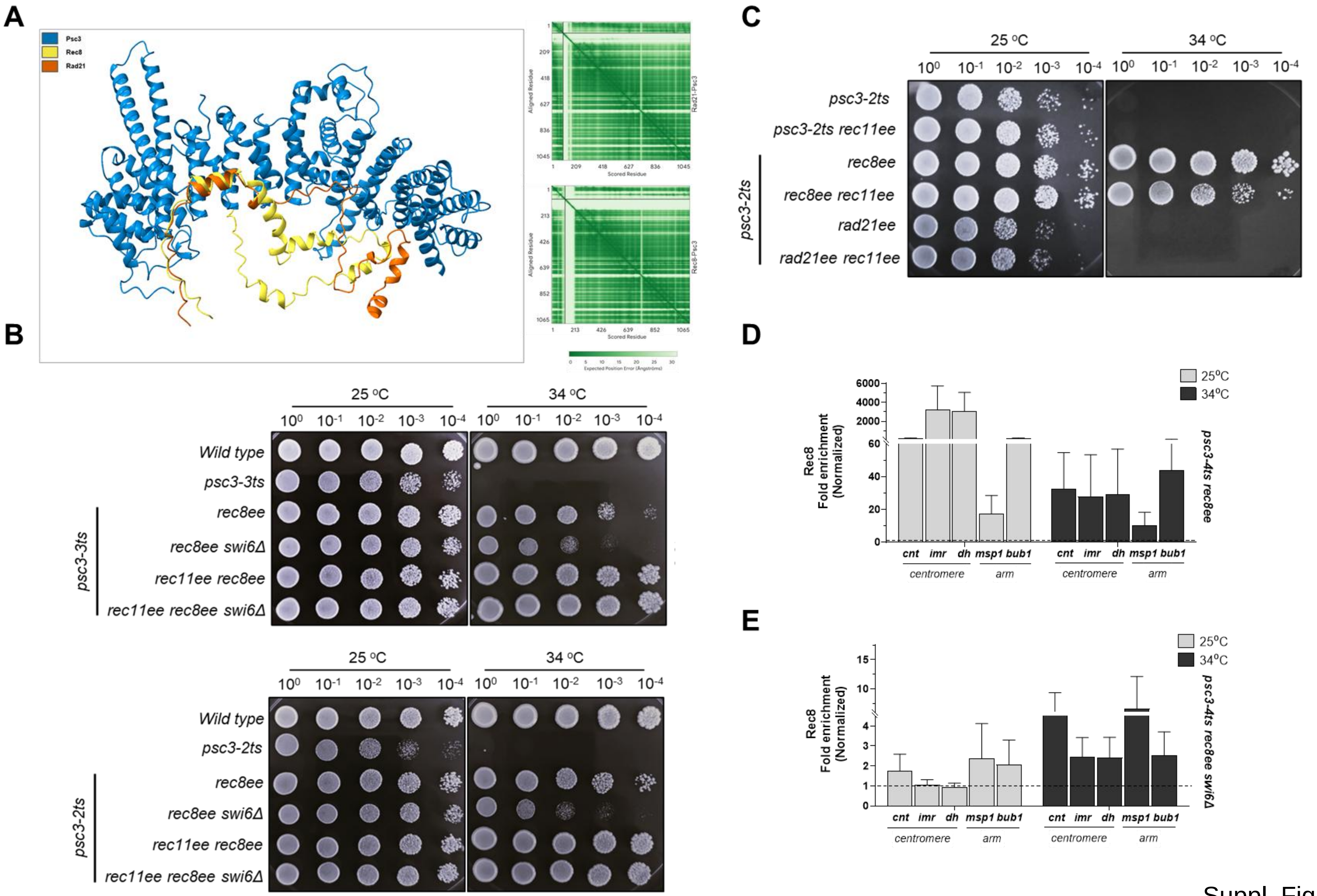
A
Psc3
Rec8
Rad21
B
25 °C
34 °C
10^0 10^-1 10^-2 10^-3 10^-4
Wild type
psc3-3ts
rec8ee
rec8ee swi6Δ
rec11ee rec8ee
rec11ee rec8ee swi6Δ
psc3-3ts
Wild type
psc3-2ts
rec8ee
rec8ee swi6Δ
rec11ee rec8ee
rec11ee rec8ee swi6Δ
psc3-2ts
C
psc3-2ts
psc3-2ts rec11ee
rec8ee
rec8ee rec11ee
rad21ee
rad21ee rec11ee
psc3-2ts
D
25°C
34°C
Rec8
Fold enrichment
(Normalized)
cnt imr dh msp1 bub1
centromere
arm
psc3-4ts rec8ee
E
psc3-4ts rec8ee swi6Δ

Suppl. Fig. 3

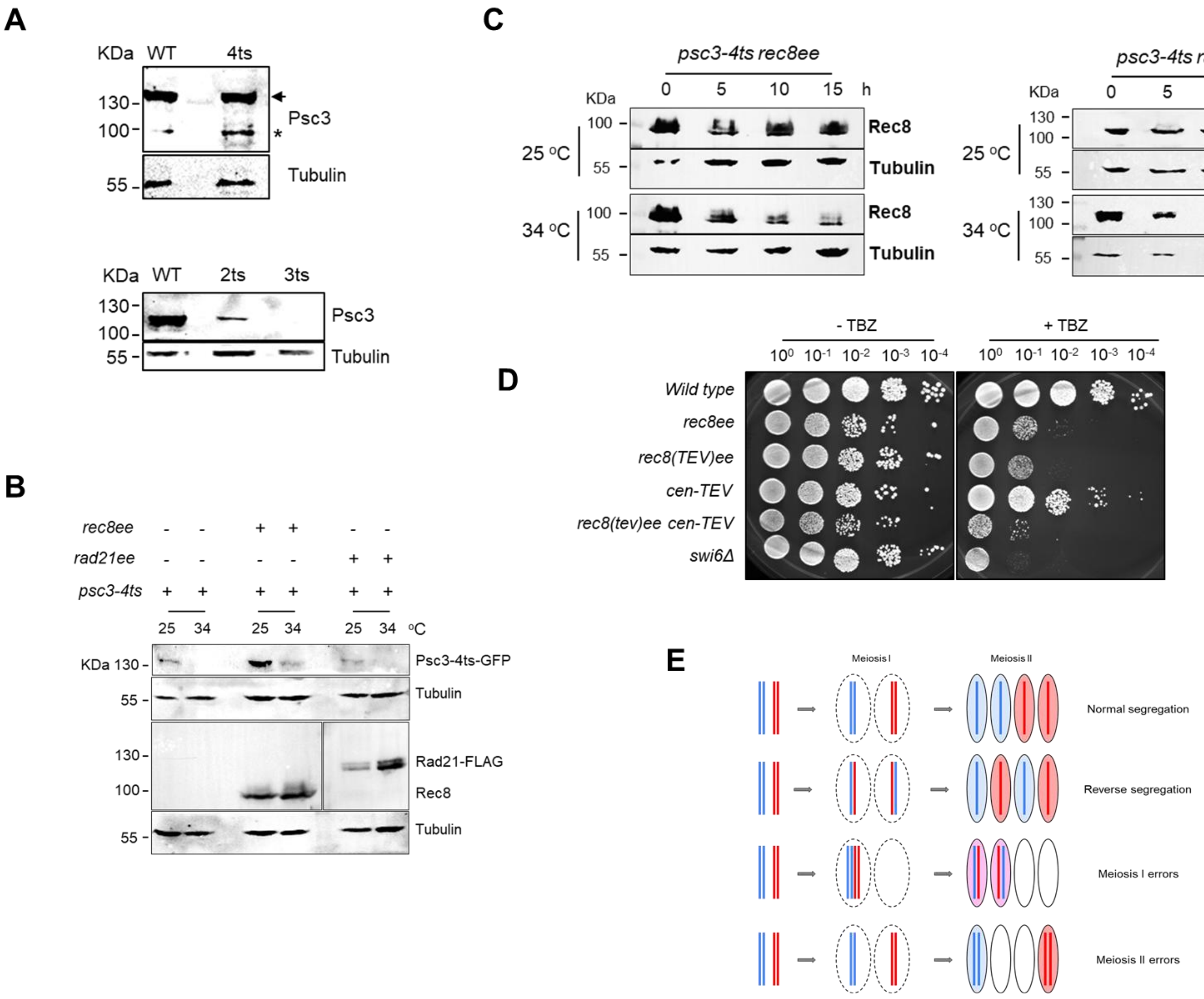
A
KDa WT 4ts
130
100
Psc3
*
55
Tubulin
KDa WT 2ts 3ts
130
100
Psc3
55
Tubulin
B
rec8ee - - + + - -
rad21ee - - - - + +
psc3-4ts + + + + + +
25 34 25 34 25 34 °C
KDa 130
Psc3-4ts-GFP
55
Tubulin
130
100
Rad21-FLAG
Rec8
55
Tubulin
C
psc3-4ts rec8ee
KDa 0 5 10 15 h
100
Rec8
25 °C
55
Tubulin
100
Rec8
34 °C
55
Tubulin
psc3-4ts rad21ee
KDa 0 5 10 15 h
130
100
Rad21
25 °C
55
Tubulin
130
100
Rad21
34 °C
55
Tubulin
D
- TBZ
+ TBZ
10^0 10^-1 10^-2 10^-3 10^-4
Wild type
rec8ee
rec8(TEV)ee
cen-TEV
rec8(tev)ee cen-TEV
swi6Δ
E
Meiosis I
Meiosis II
Normal segregation
Reverse segregation
Meiosis I errors
Meiosis II errors

Suppl. Fig. 4

**Supplementary Figure Legends**

**Supplementary Figure 1. Meiotic cohesin complex Rec8-Rec11 rescues viability upon depletion of mitotic HAWK subunit Psc3. A.** Sequencing-verified positions of point mutations in the three *psc3-ts* alleles. **B.** Spot assay for the three *psc3-ts* alleles to verify extent of temperature sensitivity at 25 $^{o}$C, 30 $^{o}$C and 34 $^{o}$C. **C.** Spot assay for *psc3-2ts* and *psc3-3ts* strains in the presence of ectopic expression (ee) of its paralog Rec11 (*rec11ee)* alone or with its meiotic partner Rec8 (*rec8ee)* at 25 $^{o}$C and 34 $^{o}$C*.* **D.** Spot assay for indicated strains at 25 $^{o}$C and 34 $^{o}$C to depict the rescue of *rad21Δ* and *psc3Δ* by ectopic expression of their meiotic paralogs, *rec8* and *rec11*, respectively*.* **E.** Growth curves for the strains in panel C at 25 $^{o}$C. The error bars represent ±SEM. **F.** Interaction between Rad21 and Rec11 in wild type cells overexpressing *rec11-HA* via co-immunoprecipitation. Rec11 was immunoprecipitated and probed with anti-Rad21 and anti-HA antibodies. M is ladder. For each experiment, atleast 3 biological repeats were performed.

**Supplementary Figure 2. Rec11-Rad21 targeting to pericentromeric heterochromatin rescues inviability due to Psc3 depletion. A.** AlphaFold predicted structures for Rec8 (yellow) and Rad21 (red) indicating the N-terminal, C-terminal domains as well as the domain for HAWK interaction (DHI). **B.** Interactions between Rec11 and the Rad21$^{DHI}$ and Rec8$^{DHI}$, predicted by AlphaFold 3. Rec11 is in grey, Rad21 in red and Rec8 in yellow. **C.** Spot assay for *psc3-2ts* strains in the presence of *rec11ee, rec11-cdee* (Rec11 fused with Swi6 chromodomain (CD))*, rec8ee* with or without *swi6Δ* at 25 $^{o}$C and 34 $^{o}$C*.* **D.** Interaction between Rec11-HA and Swi6 in wild type cells overexpressing Rec11 via co-immunoprecipitation. Swi6 was immunoprecipitated and probed with anti-HA (for Rec11) antibody. M is ladder. For each experiment, atleast 3 biological repeats were performed.

**Supplementary Figure 3. Rec8 overexpression rescues inviability of Psc3 hypomorphic mutants under restrictive conditions, unlike Rad21. A.** Interactions between Psc3 and the Domain of HAWK Interaction (DHI) in Rad21 and Rec8, predicted by AlphaFold 3. Psc3 is in cyan, Rad21 in red and Rec8 in yellow. **B.** Spot assay for *psc3-3ts* and *psc3-2ts* strains in the presence of *rec8ee* with and without *rec11ee* and/or *swi6Δ* at 25 $^{o}$C and 34 $^{o}$C*.* 1/10 dilutions were spotted on supplemented YEA media. **C.** Spot assay for *psc3-2ts* strains in the presence of *rad21ee* or *rec8ee* with and without *swi6Δ* at 25 $^{o}$C and 34 $^{o}$C*.* 1/10 dilutions were spotted on supplemented YEA media. **D-E.** ChIP qPCR for Rec8 in *psc3-4ts rec8ee* (D) and *psc3-4ts rec8ee swi6Δ* (E) at centromere and arm loci at 25 $^{o}$C and 34 $^{o}$C. Fold enrichment was calculated after normalizing with mitochondrial DNA locus as negative control. For each experiment, atleast 3 biological repeats were performed. For panels D and E, bars represent the mean of at least three independent experiments performed in triplicates; error bars=SEM.

**Supplementary Figure 4. Expression levels of Psc3-ts mutant proteins and cell viability in presence of Rec8 and Rad21. A.** Western blot to determine expression levels of all Psc3-ts mutants compared to wild-type. Tubulin is used as loading control. For Psc3-4ts, the blots were probed with anti-GFP antibody, whereas for Psc3-3ts and Psc3-2ts, anti-HA antibody was used. The wild type Psc3 used for each mutant had the respective epitope tags. "*" indicates potential cleavage band. **B.** Western blot to determine expression levels of Psc3-4ts mutant, Rec8 and Rad21 at 25 $^{o}$C and 34 $^{o}$C in the mentioned genotypes. The blots were probed with anti-GFP, anti-HA, anti-FLAG and anti-Tubulin antibodies. **C.** Time-course experiment to determine expression levels of Rec8 and Rad21 at 25 $^{o}$C and 34 $^{o}$C in non-cycling cells. The blots were probed with anti-HA (for Rec8), anti-FLAG (for overexpressed Rad21) and anti-Tubulin antibodies. M is ladder. **D.** Spot assay to assess hypersensitivity to Thiabendazole (TBZ) (0 and 7.5 µg/ml) in *wild type, rec8ee, rec8(tev)ee, cen-TEV, rec8(tev)ee cen-TEV* and *swi6Δ* strains. For each experiment, atleast 3 biological repeats were performed.
**E.** Outline for fluorescence-based tetrad assay used to determine the meiotic segregation fidelity for *wild type* and *psc3-3ts* cells, as described previously[31].

**Supplementary Table 1-** List of *Schizosaccharomyces pombe* strains used in the study.

| Strain | Genotype | Figure |
|---|---|---|
| MP1 | *wild type* | Figs. 2G, 4A, 4C, 7A and Suppl. Figs. 1B, 1D |
| MP3 | *wild type* | Figs. 1C, 2D, 7C, 7D and Suppl. Figs. 1F, 2D |
| MP21 | *ade6-52 ura4-D18 his3-D1 Padh1-rec8-3HA-ura4+ C::Padh15-rec11-hygR psc3::kanR* | Figs. 2F and Suppl. Fig. 1D |
| MP92 | *swi6::kanR* | Figs. 2F, 2G, 7A and Suppl. Fig. 4D |
| MP103 | *ade6-M216 ura4-D18 leu1::(dfp1(1-376)-6his3HA leu1+) dfp1-D1* | Figs. 7C, 7D |
| MP112 | *psc3-GFP-kanR* | Suppl. Fig. 4A |
| MP117 | *psc3-HA-[LEU2] rad21-myc ura4-D18 leu1-32 (?)* | Suppl. Fig. 4A |
| MP164 | *ade6-52 psc3-4ts-GFP-kanR* | Figs. 1A, 1B, 1D, 4A, 4B, 4C, 5C, 6A, 6B, 6C, 6D, 7C and Suppl. Figs. 1A, 1B, 4A, 4B |
| MP166 | *psc3-2ts-3HA-kanR ade6-M210* | Suppl. Figs. 1A, 1B, 1C, 1E, 2C, 3B, 3C, 4A |
| MP167 | *psc3-3ts-3HA-kanR ade6-M210* | Fig. 7A and Suppl. Figs. 1A, 1B, 1C, 1E, 3B, 4A |
| MP208 | *CEN1::his3+-PSPOG_00147-tdTomato his3-D1 ura4-D18* | Fig. 7B |
| MP238 | *CEN1::his3+-PSPOG_00147-mCerulean his3-D1 ura4-D18* | Fig. 7B |
| MP279 | *rec8D::Padh1-rad21+ -FLAG -ura4+ ade6-M210 ura4-D18* | Fig. 4C |
| MP317 | *ade6-52 ura4-D18 padh1-rec8-3HA-ura+* | Figs. 2C, 2G, 4B, 4C, 5C and Suppl. Fig. 4D |
| MP318 | *ade6-52 ura4-D18 c:: padh15-rec11-hygR* | Figs. 1A, 1B, 2G and Suppl. Figs. 1C, 1E |
| MP319 | *ade6-52 c:: padh15-rec11-hygR psc3-4ts-GFP-kanR* | Figs. 1A, 1B, 2A, 7D |
| MP320 | *ade6-M210 c::padh15-rec11-hygR psc3-2ts-3HA-kanR* | Suppl. Figs. 1C, 1E, 2C, 3C |
| MP321 | *ade6-M210 c::padh15-rec11-hygR psc3-3ts-3HA-kanR* | Suppl. Fig. 1E |
| MP361 | *ade6-M210 ura4-D18 his3-D1 rad21::ura4+ padh1-rec8-3HA-ura4+* | Suppl. Fig. 1D |
| MP362 | *ade6-52 ura4-D18 psc3-4ts-GFP-kanR Padh1-rec8-3HA-ura+ c:: Padh15-rec11-hygR* | Figs. 1A, 1B, 1F, 2A, 4A, 4B, 6E, 6F, 7D |

| | | |
|---|---|---|
| MP363 | *ade6-52 ura4-D18 psc3-3ts-3HA-kanR padh1-rec8-3HA-ura+ c:: Padh15-rec11-hygR* | Suppl. Figs. 1C, 1E, 3B |
| MP373 | *ade6-M210 ura4-D18 c::Padh15-rec11-hygR psc3-3ts-3HA-kanR* | Suppl. Fig. 1C |
| MP422 | *ade6-M210 ura4-D18 Padh1-rec8-3HA-ura4+ psc3-2ts-3HA-kanR* | Suppl. Figs. 3B, 3C |
| MP423 | *ade6-M210 ura4-D18 Padh1-rec8-3HA-ura4+ psc3-2ts-3HA-kanR c::padh15-rec11-hygR* | Suppl. Figs. 1C, 1E, 2C, 3B, 3C |
| MP424 | *ade6-52 ura4-D18 Padh1-rec8-3HA-ura4+ c::padh15-rec11-hygR* | Figs. 2F, 2G |
| MP426 | *ade6-M210 ura4-D18 psc3-2ts-3HA-kanR c::Padh15-rec11-hygR rec8D::Padh1-rad21+ -FLAG-ura4+* | Suppl. Fig. 3C |
| MP427 | *ade6-M210 ura4-D18 psc3-2ts-3HA-kanR rec8D::Padh1-rad21+ -FLAG-ura4+* | Suppl. Fig. 3C |
| MP500 | *ade6-M210 ura4-D18 c::Padh15-rec11-hygR Padh1-rec8-3HA-ura4+* | Fig. 2C |
| MP503 | *ade6-52 ura4-D18 arg1-14 psc3::kanR Padh1-rec8-3HA-ura4+ c::Padh15-rec11-hygR* | Fig. 2C |
| MP504 | *ade6-52 psc3-4ts-GFP-kanR ura4+-padh1-rec11-CD-hygR (native ura4+)* | Fig. 2A |
| MP505 | *ade6-52 ura4-D18 psc3-4ts-GFP-kanR swi6::natR ura4+-Padh1-rec11-GFP-CD-hygR* | Fig. 2A |
| MP535 | *ura4-D18 swi6::natR his3-D1 Padh1-rec8-3HA-ura4+* | Fig. 2G |
| MP537 | *ura4-D18 swi6::natR his3-D1 Padh1-rec8-3HA-ura4+ C::Padh15-rec11-hygR psc3::kanR* | Figs. 2E, 2F |
| MP539 | *ura4-D18 his3-D1 swi6::natR C::Padh15-rec11-hygR* | Fig. 2G |
| MP540 | *ura4-D18 swi6::natR his3-D1 Padh1-rec8-3HA-ura4+ C::padh15-rec11-hygR* | Fig. 2G |
| MP547 | *ade6-52 ura4-D18 his3-D1 Padh1-rec8-3HA-ura4+ psc3-4ts-GFP-kanR* | Figs. 4A, 4B, 4C, 5A, 5F, 6A, 6B, 6C, 6D, 6E, 6F, 7C and Suppl. Figs. 3D, 4B, 4C |
| MP548 | *ade6-52 ura4-D18 Padh1-rec8-3HA-ura4+ psc3-3ts-3HA-kanR* | Fig. 7A and Suppl. Fig. 3B |
| MP551 | *ade6-M26/M210 swi6::natR ura4+-Padh1-rec11-265::GFP-CD-hygR psc3-2ts-3HA-kanR* | Fig. 2B and Suppl. Fig. 2C |
| MP552 | *ade6-M26/M210 ura4-D18 ura4+-Padh1-rec11-265::GFP-CD-hygR psc3-2ts-3HA-kanR* | Fig. 2B and Suppl. Fig. 2C |

| | | |
|---|---|---|
| MP569 | *ade6-52 ura4-D18 psc3-4ts-GFP-kanR Padh1-rec8-3HA-ura4+ swi6::natR* | Figs. 4A, 5B, 6A and Suppl. Fig. 3E |
| MP571 | *ade6-M210/M26 ura4-D18 Padh1-rec8-3HA-ura4+ psc3-2ts-3HA-kanR c::Padh15-rec11-hygR swi6::natR* | Suppl. Figs. 2C, 3B |
| MP572 | *ade6-52 ura4-D18 psc3-3ts-3HA-kanR Padh1-rec8-3HA-ura4+ c:: Padh15-rec11-hygR swi6::natR* | Suppl. Fig. 3B |
| MP574 | *ade6-52 ura4-D18 psc3-4ts-GFP-kanR Padh1-rec8-3HA-ura+ swi6::natR c:: Padh15-rec11-hygR* | Figs. 2A, 4A |
| MP618 | *ade6-M210/ade6-M26 ura4-D18 psc3-2ts-HA-KanR Padh1-rec8-HA-ura4+ swi6::natR* | Suppl. Fig. 3B |
| MP620 | *ade6-52 ura4-D18 psc3-3ts-HA-kanR Padh1-rec8-HA-ura4+ swi6::natR* | Fig. 7A and Suppl. Fig. 3B |
| MP871 | *psc3-3ts-3HA-kanR CEN1::his3+-PSPOG_00147-mCerulean ura4-D18 his3-D1* | Fig. 7B |
| MP932 | *psc3-3ts-3HA-kanR CEN1::his3+-PSPOG_00147-tdTomato ura4-D18 his3-D1* | Fig. 7B |
| MP986 | *ura4-D18 ura4+-Padh1-rec11-3HA-kanR psc3-4ts-GFP-kanR* | Figs. 1D, 1E |
| MP1036 | *ade6-52 Padh1-rec8-HaloTag-hygR swi6-GFP-kanR* | Figs. 3A, 3B, 3C |
| MP1039 | *ade6-52/ade6-M216 ura4-D18 ura4+-Padh1-rec11-3HA-kanR ssl3-FLAG-kanR* | Figs. 1C, 2D, 3F and Suppl. Figs. 1F, 2D |
| MP1101 | *ade6-M210 ura4-D18 his3-D1 rec8D::Padh1-rad21+ -FLAG-ura4+ psc3-4ts-GFP-kanR* | Figs. 4C, 6B, 6C, 6D and Suppl. Figs. 4B, 4C |
| MP1278 | *ade6-52/ade6-M216 ura4-D18 ura4+-Padh1-rec11_lng-lnk_HaloTag-HygR swi6-GFP-kanR* | Figs. 3A, 3D, 3E |
| MP1321 | *ade6-M216 leu1- ura4-D18 rec8::Padh1-rec8(TEV)-3HA-ura4+* | Fig. 5C and Suppl. Fig. 4D |
| MP1322 | *ade6-M216 leu1- lys1:: Padh81-cen-TEV-hygR* | Suppl. Fig. 4D |
| MP1323 | *ade6-M216 leu1- lys1::Padh81-cen-TEV-hygR rec8::rec8(TEV)* | Fig. 5C and Suppl. Fig. 4D |
| MP1336 | *ade6-52/ade6-M216 (?) ura4-D18 Padh1-rec8-HaloTag-hygR C::Padh15-rec11-hygR swi6-GFP-kanR* | Figs. 3B, 3C |
| MP1338 | *ade6-52/ade6-M216 (?) ura4-D18 Padh1-rec8-HaloTag-hygR C::Padh15-rec11-hygR swi6-GFP-kanR psc3::natR* | Figs. 3B, 3C |
| MP1377 | *ade6-52/ade6-M216 ura4-D18 leu1::(dfp1(1-376)-6his3HA leu1+) dfp1-* | Fig. 7D |

| | | |
|---|---|---|
| | *D1 c::Padh15-rec11-hygR psc3-4ts-GFP-kanR Padh1-rec8-3HA-ura4+* | |
| MP1412 | *ade6-52/ade6-M216 ura4-D18 Padh1-rec8-3HA-ura4+ ura4+-Padh1-rec11_lng-lnk_HaloTag-HygR swi6-GFP-kanR* | Figs. 3D, 3E |
| MP1413 | *ura4-D18 Padh1-rec8-HaloTag-hygR psc3-FLAG3-NatR* | Fig. 2D |
| MP1423 | *ade6-52/ade6-M216 ura4-D18 leu1-32 rec8::Padh1-rec8(TEV)-3HA-ura4+ lys1:: Padh81-cen-TEV-hygR* | Suppl. Fig. 4D |
| MP1424 | *ade6-52/ade6-M216 ura4-D18 his3-D1 leu1-32 rec8::Padh1-rec8(TEV)-3HA-ura4+ lys1:: Padh81-cen-TEV-hygR psc3-4ts-GFP-kanR* | Figs. 5C, 5D, 5E, 5F |
| MP1425 | *ura4-D18 Padh1-rec8-HaloTag-hygR ura4+-Padh1-rec11-3HA-kanR psc3-FLAG3-natR* | Figs. 2D, 3F |
| MP1447 | *ade6-52/ade6-M216 ura4-D18 leu1::(dfp1(1-376)-6his3HA leu1+) dfp1-D1 Padh1-rec8-3HA-ura4+ psc3-4ts-GFP-kanR* | Fig. 7C |

**Supplementary Table 2-** List of oligonucleotides used in the study.

| Oligo # | Sequence | Used for |
|---|---|---|
| MO53 | AGAAAAAGGAGTTGGTGGGTTCTCAAAAGGGGTCAGGGAAAAGCCCGAAATTCCGAGTAAAAACGCGTTCTTATTCATAAGGAATGCCATGTCAGATTTG | *Padh1-rec11* allele generation |
| MO54 | TTTTCAAGCTCCACGCTTTCTAATTCAGTAAGGACTTCATTGAGATGACTGCTTGCATCTGATTCAAATTCGAATCTCATGGATCCGCTAGCGGAATTCT | |
| MO341 | TAGTGTCTCTTTAAACCTAAAAAGCACCGATGCCATTATGGGTTCCGAAATCGGTACTG | *rec8_HaloTag* allele generation |
| MO342 | CGTTTTAAAAATTTTAAGTTTAACATGTGAAAAGTTTCAGGATGGCGGCGTTAGTATCG | |
| MO343 | TGCATTTTTAGTTAAACAGGATAAACCATACTCTGAAATTAGTGTCTCTTTAAACCTAA | |
| MO344 | TTTGACAACTTCAACAAAGGGTTTAATTCCCAAATTCAACGTTTTAAAAATTTTAAGTT | |
| MO468 | CAAGCGGCCGCCGCTGCTGCCGCTGCAGCTGCTGCAGCTGCGATGGGTTCCGAAATCGG | *rec11_HaloTag* allele generation |
| MO375 | TTTCCCTCGGGGTTTAATTAGTGTTAAAATTTCGTTTTAGGATGGCGGCGTTAGTATCG | |
| MO471 | GGTATCCCAATCGTGCTAGAGATCCTGGAACACTGAGTCATTTAGTGAAAGGTTTAAAAGAAACAGCTGACCATTTATCACAAGCGGCCGCCGCTGCTGC | |
| MO377 | CCTCAATAAGAAAGTTAGCACATGTGCTTTCTTATCTCTTTTCCCTCGGGGTTTAATTA | |
| MCO1 | CAACTTACATCAGCATACTGG | ChIP - $cnt_{All}$ |
| MCO2 | CGAAAGATGGTCAATTGCTT | |
| MCO5 | AAAGCAAAAACCGAGTTTGG | ChIP - *imrIII* |
| MCO6 | AAATTTCATGAACGCTGACG | |
| MCO25 | GGAATAGCATACCGTCAAGTCGTTAGTTG | ChIP - $dh_{All}$ |
| MCO26 | GGTCAACGCACGCCTAAACTAGC | |
| MCO33 | GAAATCTAGTCGAGGTCAAG | ChIP – *msp1* (Chr. II) |
| MCO34 | CTTCCAAGTACTGCAAAACC | |
| MCO37 | CGTCCAATTTAGGCCAT | ChIP – *bub1* (Chr. III) |
| MCO38 | GTCTTAACCGTCTCCCTG | |
| MCO51 | ACCAGTACACGAACACGCATT | ChIP – *MitoNeg* |
| MCO52 | ATCCTTCAATCTCCCTCTCCA | |
